# A nucleosome switch primes Hepatitis B Virus infection

**DOI:** 10.1101/2023.03.03.531011

**Authors:** Nicholas A. Prescott, Andrés Mansisidor, Yaron Bram, Tracy Biaco, Justin Rendleman, Sarah C. Faulkner, Abigail A. Lemmon, Christine Lim, Pierre-Jacques Hamard, Richard P. Koche, Viviana I. Risca, Robert E. Schwartz, Yael David

## Abstract

Chronic hepatitis B virus (HBV) infection is an incurable global health threat responsible for causing liver disease and hepatocellular carcinoma. During the genesis of infection, HBV establishes an independent minichromosome consisting of the viral covalently closed circular DNA (cccDNA) genome and host histones. The viral X gene must be expressed immediately upon infection to induce degradation of the host silencing factor, Smc5/6. However, the relationship between cccDNA chromatinization and X gene transcription remains poorly understood. Establishing a reconstituted viral minichromosome platform, we found that nucleosome occupancy in cccDNA drives X transcription. We corroborated these findings in cells and further showed that the chromatin destabilizing molecule CBL137 inhibits X transcription and HBV infection in hepatocytes. Our results shed light on a long-standing paradox and represent a potential new therapeutic avenue for the treatment of chronic HBV infection.

## Introduction

Over 325 million people worldwide are chronically infected by Hepatitis B Virus (HBV), leading to almost one million deaths annually despite the existence of an effective vaccine.^1^ Chronic infection with this incurable virus leaves patients at risk for advanced liver disease, and HBV is estimated to cause nearly half of all cases of hepatocellular carcinoma.^2^ Infection begins with viral entry into hepatocytes via the sodium taurocholate co-transporting polypeptide (NTCP) bile acid receptor.^3^ Next, the capsid is shuttled to the nuclear pore, where it disassembles to release the 3.2 kb partially double-stranded, relaxed circular DNA (rcDNA) HBV genome into the nucleus.^4,5^ Following nuclear import, rcDNA is repaired into fully double-stranded, covalently closed circular DNA (cccDNA) by host cell lagging strand synthesis machinery.^3,6,7^ Host histones are rapidly deposited onto cccDNA, which then serves as the primary template for viral transcription.^8–10^ Presently, long-term treatment with oral nucleos(t)ide analogs or short-term treatment with interferon-alpha injections remains the standard of care to halt HBV replication, but these fall short of eradicating cccDNA in infected hepatocytes, allowing the viral minichromosome to persist and sustain chronic infection.^11^

Even with long-term antiviral treatment, basal levels of the HBV oncogenic X protein (HBx) remain present in hepatocytes to promote host genome instability and disease progression.^1,2,12^ HBx induces the degradation of the host Smc5/6 complex, which otherwise would transcriptionally silence cccDNA.^13,14^ However, HBx is absent from the virion and must be expressed *de novo* from cccDNA in freshly infected hepatocytes. The X gene, which encodes HBx, is the earliest-expressed viral gene, with its transcript most abundant in the first 24 hours after HBV infection.^15,16^ Although some reports identified X mRNA circulating in the plasma of infected patients, it has yet to be proven whether it is truly packaged alongside HBV rcDNA into infectious virions.^15,17^ This seemingly paradoxical relationship to cccDNA transcription during the earliest stages of infection necessitates further study of the regulation of X transcription.

Epigenetic mechanisms have been suggested to play a role in the overall regulation of viral transcription, as cccDNA is decorated predominantly with active transcription-associated histone post-translational modifications (PTMs).^8,18–21^ However, the mechanistic interplay between cccDNA chromatin state and viral transcription remains poorly characterized despite its potential importance in the establishment of active infection.

This gap in understanding arises primarily because traditional genetic and cell biology approaches lack the temporal and biochemical resolution necessary to characterize critical early intermediates in HBV infection. To surmount these difficulties, we developed a method to generate recombinant, chromatinized cccDNA in a chemically defined system, allowing us to characterize its biophysical properties and map nucleosome positioning on the minichromosome. We applied biochemical and cellular techniques to further demonstrate that nucleosome occupancy flanking the X gene transcription start site (TSS) may enhance its expression *in vitro* and in cells. Finally, we demonstrated that small molecule-mediated disruption of nucleosome integrity in cccDNA is sufficient to inhibit X gene transcription and antagonize viral infection in cultured cells and primary human hepatocytes.

## Results

### Recombinant cccDNA robustly assembles into minichromosomes *in vitro*

Previous approaches to generate cccDNA *in vitro* have largely relied on recombination-based methods, which leave undesirable exogenous DNA “scars” in the viral genome, or multi-step preparations of rcDNA for repair by recombinant proteins with comparatively low yields.^6,22,23^ Moreover, to the best of our knowledge, the only prior study of chromatinized cccDNA *in vitro* used *Xenopus laevis* egg extracts for chromatin assembly, which are transcriptionally silent and yield chemically heterogeneous chromatin fibers suboptimal for detailed biochemical interrogation.^10,24^ To overcome these limitations, we first generated a large-scale recombinant double-stranded linear HBV DNA (dslDNA). The dslDNA was then subjected to intramolecular enzymatic ligation to produce scarless cccDNA, as has been done previously to generate DNA minicircles for studies of chromatin topology (Fig. 1A, Fig. S1A).^25^ To validate the infectious capacity of the resulting recombinant cccDNA, we transfected it into HepG2 cells and assayed for hallmarks of HBV infection. Intracellular viral RNA and secreted viral DNA were readily detected for up to two weeks post-transfection (Fig. 1B). Likewise, HBV surface and secreted antigens (HBsAg and HBeAg, respectively) were abundant in the supernatant of transfected cells (Fig. S1B). Finally, virions collected from the supernatant of cccDNA-transfected cells proved capable of inducing a secondary infection when added to HepG2 cells constitutively expressing the NTCP entry receptor (HepG2-NTCP), as validated by immunofluorescence to detect intracellular viral capsid protein (HBcAg) (Fig S1C). Together, these results validate our approach to generate cccDNA without recombination scars at sufficiently high yield for biochemical and cellular studies.

**Figure 1.**
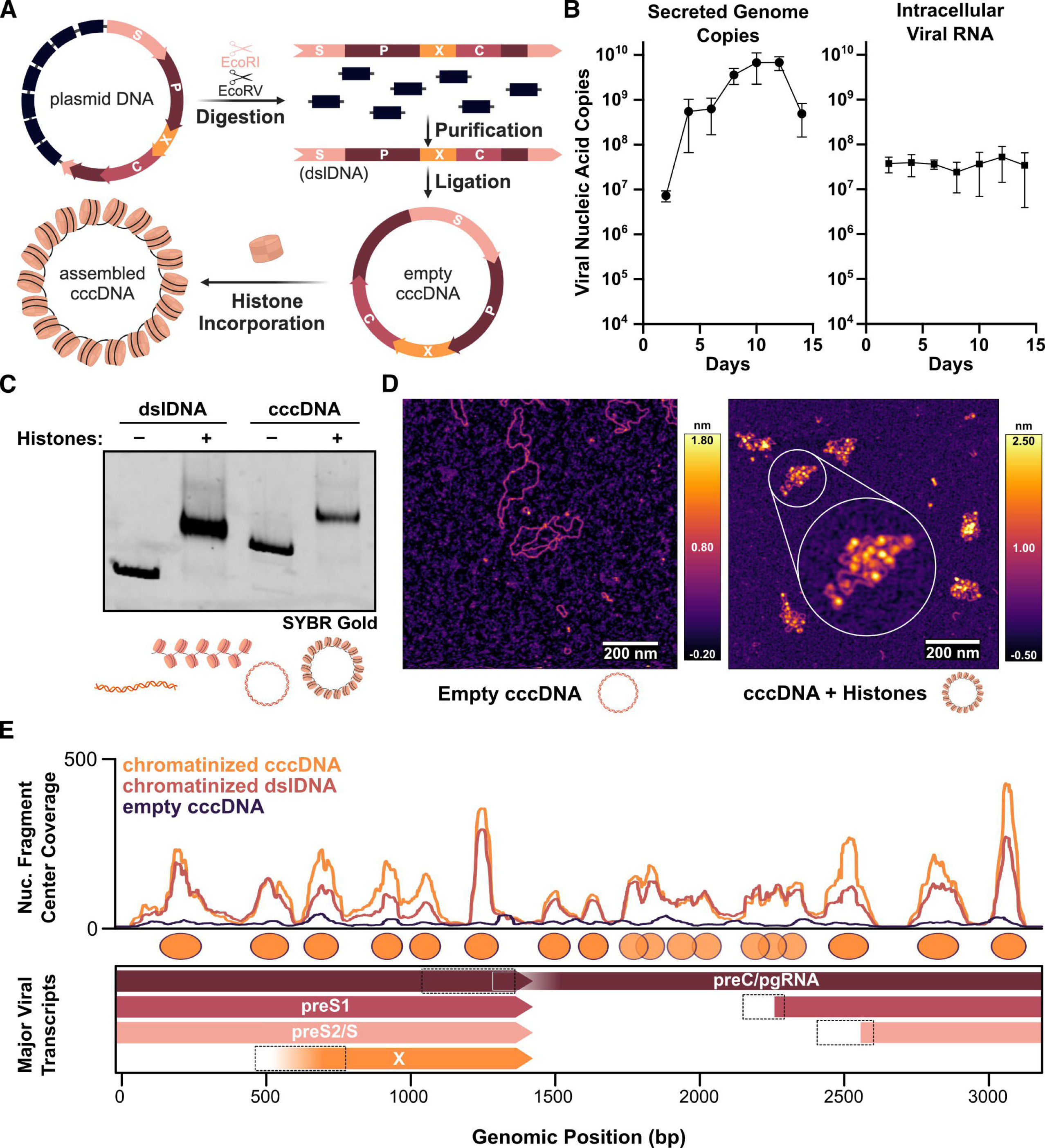
Recombinant cccDNA robustly assembles into minichromosomes *in vitro*. (A) Scheme illustrating minichromosome reconstitution approach. (B) Quantification of secreted HBV DNA (left) and intracellular total HBV RNA (right) following transfection of recombinant cccDNA into HepG2 cells. Data represent means ± SD of 3 biological replicates. (C) Electrophoretic mobility shift of linear (dslDNA) and circular (cccDNA) HBV DNA upon incorporation of recombinant human histone octamers. (D) Representative atomic force micrographs of empty (left) or chromatinized (right) cccDNA. (E) Coverage of centers of mononucleosome-length (140-200 bp) DNA fragments arising from MNase digestion of chromatinized cccDNA (orange), dslDNA (red), or empty cccDNA (navy) aligned to the HBV genome and smoothed with a 50 bp window chosen to show partially overlapping nucleosome occupancy peaks that may indicate underlying heterogeneity in nucleosome positioning, overlaid above a schematic of the four major HBV transcripts. Dotted boxes indicate promoter regions for each transcript, in genomic coordinate order: enhancer I/X promoter, enhancer II/basal core promoter, S1 promoter, S2 promoter.

We next tested our recombinant cccDNA as a substrate for *in vitro* chromatin assembly. To do so, we combined purified canonical human histone octamers with the viral DNA and used salt-gradient dialysis to assemble chromatin fibers. Histone octamer incorporation into both dslDNA and cccDNA was first validated by electrophoretic mobility shift assay (EMSA) of the intact chromatin fibers (Fig. 1C). We subsequently visualized histone-free and chromatinized cccDNA using atomic force microscopy (AFM) to illustrate the robust assembly of viral minichromosomes (Fig 1D). We next sought to assess the basic biophysical characteristics of the reconstituted HBV chromatin, which we compared to similarly-sized nucleosome arrays comprised of 18 repeats of the Widom 601 strong nucleosome-positioning sequence used to model chromatin arrays.^26^ The corresponding linear and circular constructs, Lin601^18^ and Cir601^18^, respectively, matched both the length and predicted number of nucleosomes in the HBV genome, and similarly assembled into chromatin fibers (Fig. S1D-F).^27^ We next quantitated a basal chromatin compaction metric for nucleosome arrays of all four species (dslDNA, cccDNA, Lin601^18^ and Cir601^18^) by measuring the ratio between the volume and occupied surface area of individual particles in AFM micrographs (Fig. S1G). To orthogonally compare the compaction of the fibers, we also measured their propensity to precipitate in a magnesium-dependent self-association assay (Fig. S1H).^28^ Our results indicate that *in vitro* assembled HBV and 601 chromatin arrays display similar degrees of compaction.

Finally, we sought to determine whether the reconstituted viral chromatin exhibits any DNA-intrinsic nucleosome organization, or if nucleosome positioning is stochastic *in vitro*. Micrococcal nuclease digestion followed by sequencing (MNase-seq) of both cccDNA and dslDNA chromatin arrays revealed a comparable chromatin architecture between the two species, suggesting a degree of sequence-intrinsic nucleosome positioning by the HBV genome seemingly unaffected by template circularization (Fig. 1E, S1I). Both species displayed several tightly positioned nucleosome-sized DNA fragments (140-200 bp), as well as some regions without strong nucleosome-sized fragment peaks, likely due to more heterogeneous nucleosome positioning. Additionally, we used an orthogonal chaperone-assisted assembly method to chromatinize dslDNA and assess nucleosome positioning therein. The *S. cerevisiae* histone chaperone Nap1, alone or together with the *Drosophila* ACF chromatin remodeler complex, was used to assemble dslDNA chromatin fibers as others previously have applied with 601 templates.^29–31^ Following assembly, we subjected both samples to MNase-seq. We found that Nap1 and Nap1-ACF assembled fibers exhibited similar patterns of nucleosome positioning, with a slightly greater density of well-positioned nucleosomes relative to the salt dialysis-assembled fibers and more accessibility, enhancing MNase digestion down to mononucleosomes-sized fragments (Fig. S1J-K). Some, but not all, nucleosome peaks remained consistent between the two approaches, suggesting that the HBV genome may exhibit a partial intrinsic propensity to position nucleosomes. Altogether, our *in vitro* characterization efforts have validated our approach to reconstitute cccDNA minichromosomes as a robust platform to enable further mechanistic investigation of HBV chromatin.

### Chromatinization of reconstituted cccDNA enhances transcription of the viral X gene

To examine the role of chromatin in HBV transcription, we first tested the capacity of recombinant cccDNA to serve as a template for *in vitro* transcription. We incubated histone-free or chromatinized cccDNA with transcriptionally competent HeLa cell nuclear extract and then quantified total transcribed viral RNA by RT-qPCR. Unexpectedly, we detected a significant increase in the transcriptional output of the chromatinized template compared to empty DNA (Fig. S2A). These results contradicted conventional findings that nucleosomes are obstacles to transcription rather than activators.^32,33^ To ensure that the cell-type origin of the nuclear extract used for *in vitro* transcription had no effect on our observations, we repeated the assay with nuclear extracts prepared from HepG2 cells.^34^ Both HeLa and HepG2 extracts were free of histones but enriched for soluble nuclear proteins such as RNA Polymerase II and the FACT complex, among others (Fig. S2B). Consistent with our results in HeLa extract, chromatinized cccDNA displayed an enhanced transcriptional output compared to the unchromatinized template in HepG2 nuclear extract (Fig. 2A). To rule out the possibility that the increased transcriptional signal was due to degradation of the histone-free template, we confirmed that neither extract digested DNA (Fig. S2C). We also validated our analyses using full-length 5’ RACE on RNA extracted from cccDNA-transfected HepG2 cells and *in vitro* transcription reactions on chromatinized cccDNA. Although we observed differences in transcript abundance and heterogeneity in the X transcript length *in vitro*, both transcriptomes exhibited each of the major viral transcripts: the 3.5 kb pregenomic RNA (pgRNA) and preC transcripts (encoding the HBeAg, capsid, and polymerase proteins), the 2.4 kb preS1 transcript (encoding the longest HBsAg isoform), the 2.1 kb preS2 transcript (encoding two shorter HBsAg isoforms), and the 0.7-0.9 kb X transcript (Fig. S2D).

**Figure 2.**
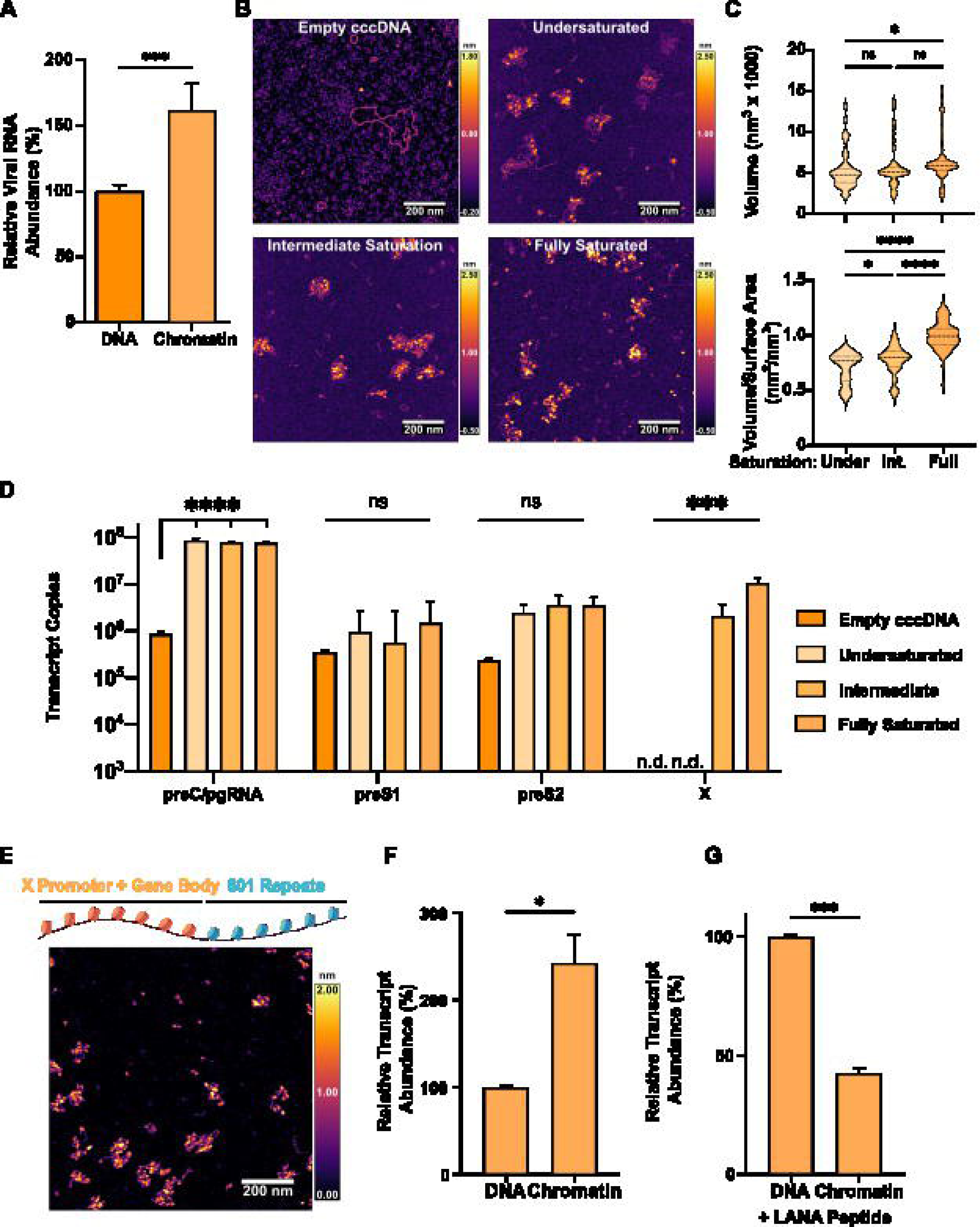
Chromatinization of reconstituted cccDNA impacts viral transcription. (A) Quantification of relative abundance of HBV RNA produced via *in vitro* transcription (IVT) of empty or chromatinized cccDNA in HepG2 nuclear extract. (B) Representative AFM micrographs of empty, undersaturated, intermediately, or fully saturated cccDNA minichromosomes. (C) Quantification of chromatin fiber volume (top) and volume-surface area ratio, a proxy for compaction, (bottom) for n = 49 (undersaturated), 70 (intermediate), or 96 (saturated) chromatin fibers pooled from multiple sample preparations on multiple days. (D) Quantification of transcript abundance for each of the four major HBV transcript following IVT of empty, undersaturated, intermediately, or fully saturated cccDNA species in HeLa nuclear extract. (E) Schematic depiction (top) and representative atomic force micrograph (bottom) of chimeric X-601 chromatin array comprised of the X promoter and gene body fused upstream of six 601 repeats. (F) Quantification of X transcript produced following IVT of empty or chromatinized chimeric X-601 template DNA, relative to an internal control, in HepG2 nuclear extract. (G) IVT as in E, with 1 µM synthetic LANA peptide added to all reactions. Data represent means of 3 replicates ± SD and were analyzed by Welch’s t-test (A, C, F, G) and 2-way ANOVA with Dunnett’s multiple comparison T3 test (D). *P < 0.05, **P < 0.01, ***P < 0.001, ****P <0.0001.

In order to monitor individual transcripts, rather than global HBV transcription, we turned to a targeted RT-qPCR assay. This assay enables the absolute quantitation of the four major viral transcripts by using different primer pairs to measure the abundance of both overlapping and non-overlapping regions of the viral transcripts and subtracting the signals from one to another.^35^ As substrates, we generated a series of cccDNA minichromosomes with increasing levels of histone saturation – under-, intermediately, or fully saturated – by altering the stoichiometry of octamer to DNA during chromatin assembly. The three species were qualitatively and quantitatively distinct from one another when visualized by AFM (Fig. 2B). As expected, increasing histone octamer saturation correlated with greater chromatin compaction (Fig. 2C). We next tested whether varying histone saturation in cccDNA influenced *in vitro* transcription. Interestingly, not all viral transcripts respond equally to the presence of chromatin (Fig. 2D, S2E). The preS1 and preS2 transcripts’ expression was not significantly changed by template chromatinization, whereas preC/pgRNA expression significantly increased with chromatinization although it was still transcribed from empty DNA. Notably, there was no significant difference between the preC/pgRNA output of templates from the three different saturation levels. Strikingly, we observed a switch-like behavior for the X transcript: empty and undersaturated cccDNA did not detectably produce X transcript, whereas intermediately and fully saturated templates did (Fig. 2D). Moreover, this increased expression of X did not come at the expense of other transcripts: although the relative proportions of non-X transcripts decreased with greater template histone saturation, the absolute abundance of each transcript was steady or increased with greater saturation (Fig. S2E).

The notion that the X gene, known to be a critical early transcript during viral infection, could require nucleosome occupancy for expression motivated us to further explore the relationship between its chromatinization and transcription. To do so, we generated a simplified *in vitro* transcription template termed X-601, comprised of the X promoter and gene body fused upstream of six repeats of the 601 nucleosome-positioning sequence to increase the size of the reconstituted chromatin fibers and mimic the remainder of the viral minichromosome (Fig. 2E). Consistent with our results on intact cccDNA, in both HeLa and HepG2 extracts we detected a significant increase in X transcription from chromatinized templates, relative to an internal control (Fig. 2F, S2F). To further probe whether histone-dependent interactions are involved in this enhancement, we synthesized a peptide derived from the Kaposi’s sarcoma virus latency-associated nuclear antigen (LANA) (Fig. S2G), which binds the nucleosome acidic patch and can compete off histone-dependent nucleosome interactions, for example by chromatin remodelers or the transcription preinitiation complex (PIC)-Mediator supercomplex.^36–38^ Indeed, addition of LANA peptide to *in vitro* transcription reactions fully ablated the enhancement of X-601 transcription by chromatinization, instead resulting in an approximate 60 % decrease in the transcription of chromatinized X-601 templates compared to empty DNA (Fig. 2G, S2H). Together these results suggest that chromatinization of the X promoter enhances its transcriptional capacity *in vitro*, facilitated by acidic patch-dependent interactions.

### Early X gene transcription is linked to the established cccDNA nucleosome landscape

To better understand the relationship between cccDNA chromatinization and X gene transcription, we first set out to assess nucleosome occupancy patterns via MNase-seq in the three increasingly histone-saturated species of cccDNA used as *in vitro* transcription templates. All samples yielded peaks of mononucleosome-sized DNA fragment coverage at similar loci across the viral genome, although the signal strength for certain peaks varied slightly between samples (Fig. 3A, S3A). For example, the well-positioned nucleosome with the strongest occupancy signal across all samples was centered at ∼1250 bp near the 3’ end of the HBx open reading frame (ORF) and just upstream of the preC/pgRNA TSS. Surprisingly, we could not detect clear differences in the nucleosome positioning patterns between the intermediately and fully saturated samples from which X was transcribed and the undersaturated samples from which it was absent. All three samples exhibited well-positioned nucleosomes in the X promoter, possibly flanking the X gene TSS, centered at genomic coordinates ∼500 and ∼700 bp (Fig. 3A, dotted boxes on transcript map represent promoters), reminiscent of the -1 and +1 nucleosomes observed at active TSSs throughout eukaryotic genomes.^39,40^ We speculate that the lack of observed differences, particularly considering the conformational differences observed via AFM, may be due to limitations of MNase-seq as a bulk method for measuring nucleosome occupancy. A large fraction of HBV genome copies or regions without nucleosomes are likely to be completely digested and lost from the library. To benchmark our observed *in vitro* nucleosome positioning patterns against those of cccDNA in cells we performed MNase-seq on HepAD38 and HepG2.2.15 cells, which both support cccDNA production from chromosomally integrated copies of the HBV genome.^41,42^ We observed a redistribution of most well-defined nucleosome peaks on cccDNA originating from these cells compared to reconstituted viral chromosomes, likely indicative of nucleosome positioning by active remodeling in cells (Fig. 3B, S3B). Nevertheless, the putative -1/+1 nucleosome occupancy around the X promoter at genomic coordinates ∼500 and ∼700 bp, respectively, not only remained, but was strengthened. Additionally, even after optimization of enzyme concentrations, we observed a higher sensitivity of HBV genomic chromatin to MNase digestion compared to bulk host chromatin, potentially suggesting a more accessible conformation (Fig. S3C). To probe the occupancy of both RNA Polymerase II (RNAPII) and the promoter-associated histone PTM H3K4me3 on cccDNA, we performed chromatin immunoprecipitation coupled to sequencing (ChIP-seq) on HepAD38 cells. Consistent with past reports, both H3K4me3 and RNAPII were highly enriched at TSSs across the human genome, as well as broadly distributed along the viral genome, with the X gene showing some enrichment (Fig. S3D-E).

**Figure 3.**
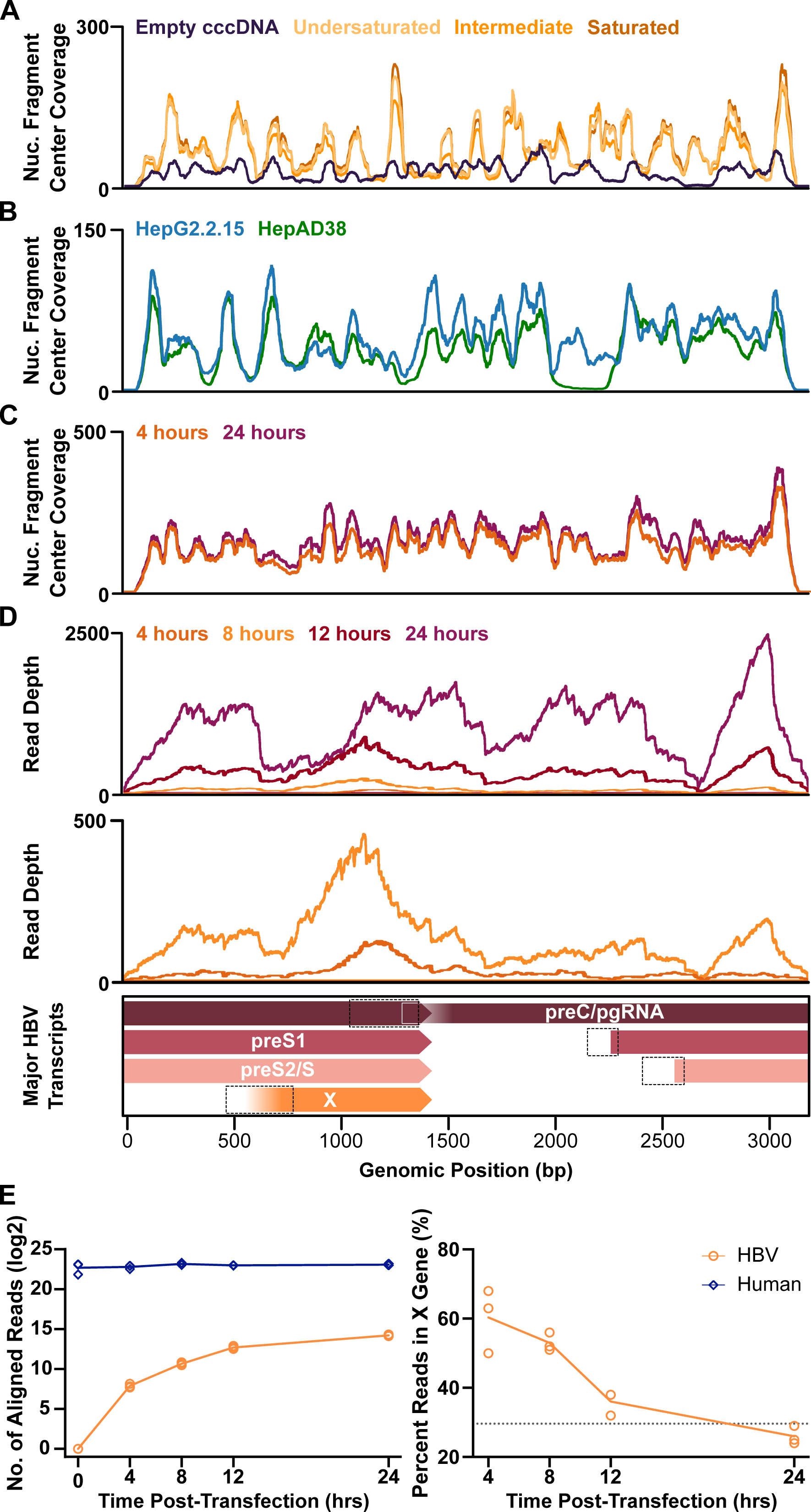
Early X gene transcription is linked to the established cccDNA nucleosome landscape. (A) Mononucleosome-sized fragment center coverage over the HBV genome following digestion of cccDNA chromatin arrays prepared as in Fig. 2B with different levels of histone octamer saturation (B) Coverage of the HBV genome by centers of unique molecular identifier (UMI)-deduplicated mononucleosome-sized fragments from MNase-seq of HepG2.2.15 (blue) and HepAD38 (green) cells. (C) Coverage of the HBV genome by centers of unique molecular identifier (UMI)-deduplicated mononucleosome-sized fragments from MNase-seq 4 (orange) and 24 hours (plum) after transfection of HEK293T cells with recombinant cccDNA. (D) HBV genomic coordinates of RNA-seq reads mapped at all time points measured up to 24 hours after cccDNA transfection into HEK 293T cells (top) and zoomed in on 4 and 8 hour time points (bottom). Dotted boxes are as in Fig. 1. (E) Quantification of total number of aligned reads to HBV or human genomes at various RNA-seq time points (left), and of the fraction of HBV-aligned reads mapping to the HBx ORF (right) for 3 biological replicates.

We next aimed to determine whether nucleosome positioning in cccDNA is altered during early time points after its establishment in nuclei. To maximize temporal control and preclude signal from residual viral DNA from infection and/or replication, we transfected unchromatinized cccDNA into HEK293T cells, which were then subject to MNase-seq after 4 or 24 hours (Fig. 3C, Fig. S3F). At both time points nucleosome-sized fragments were detectable across the viral genome, but a larger number of weak nucleosome peaks replaced the strongly positioned nucleosomes found in cells. For certain loci, occupancy patterns largely resembled those of HepAD38 and HepG2.2.15 cells, as in the region spanning approximately 2400-3200 bp of the viral genome. Other regions displayed more disorganized architecture, such as around the X promoter, where rather than clear -1/+1 nucleosomes, overlapping nucleosome peaks centered at ∼500 and ∼550 bp are visible, while a potential +1 nucleosome peak centered on ∼700 bp is weakly detectable. These observations likely reflect a greater dynamicity in the chromatin organization of the transfected cccDNA at these early times post-transfection, in a context where not all viral genome copies are required to be transcriptionally competent.

Having surveyed the nucleosome organization of cccDNA, we proceeded to assess the kinetics of HBV transcription during the early stages of infection. We similarly transfected HEK293T cells with recombinant cccDNA and sequenced poly(A)-enriched RNA at several intervals in the first 24-hours post-transfection. Concordant with literature precedent for HBV infection, modest human gene expression changes were detected in transfected cells compared to non-transfected cells at any time point (Fig. S3G).^15^ At all times post-transfection, sequencing reads readily aligned to the human genome, whereas we detected an increasing abundance of HBV reads over time, suggesting active transcription of transfected cccDNA (Fig. 3E).

Sequencing reads mapped along the entire HBV genome as early as 12 hours post-transfection, but at the two earliest time points (4 and 8 hours) reads formed a peak centered on the X ORF, consistent with reports that X is the first viral transcript to accumulate in newly infected cells (Fig. 3D).^16,17^ Although short-read sequencing techniques cannot readily discriminate between overlapping transcripts, we nevertheless attempted to quantify the fraction of HBV reads mapping to the HBx ORF, which comprises only 28 % of the viral genome. Indeed, this quantification showed that at the earliest time points upwards of 60 % of sequencing reads map to HBx (Fig. 3E). Together these data suggest a correlation between early cccDNA chromatinization and X transcription, which both occur during the first hours of infection.

### Nucleosome destabilizing drugs disrupt cccDNA integrity and inhibit viral transcription

Based on our results suggesting that chromatinization of cccDNA enhances X transcription, we hypothesized that disrupting chromatin assembly might inhibit viral infection. Nucleosome assembly in cells relies on an intricate network of histone chaperones, ATP-dependent chromatin remodelers, replisome components, and other structural proteins within the nucleus.^43^ Although relatively few pharmacological options are available to specifically inhibit these factors, we acquired and tested five such small molecules: the BAF complex inhibitors PFI-3 and BD98, the indirect FACT complex inhibitor CBL137, and the Cdc7 kinase inhibitors TAK-931 and PHA-767491 (as Cdc7 kinase activity has been shown to activate the (H3-H4)2 chaperone CAF-1).^44–49^ Following a 24-hour pre-treatment with each inhibitor, HEK 293T cells were transfected with recombinant cccDNA and drug treatment was continued until harvesting cells 4, 8, or 24 hours later for analysis. None of the doses we used impacted cell viability for up to 48 hours of treatment (Fig. S4A, Table S1), and although four of the molecules tested showed no substantial repressive effect, CBL137 was able to significantly inhibit viral transcription at all three time points (Fig. 4A). This effect proved to be dose-dependent, with CBL137 displaying an EC50 of approximately 135 nM at 24 hours post-transfection (Fig. 4B), implicating CBL137 as a potent inhibitor of HBV transcription.

**Figure 4.**
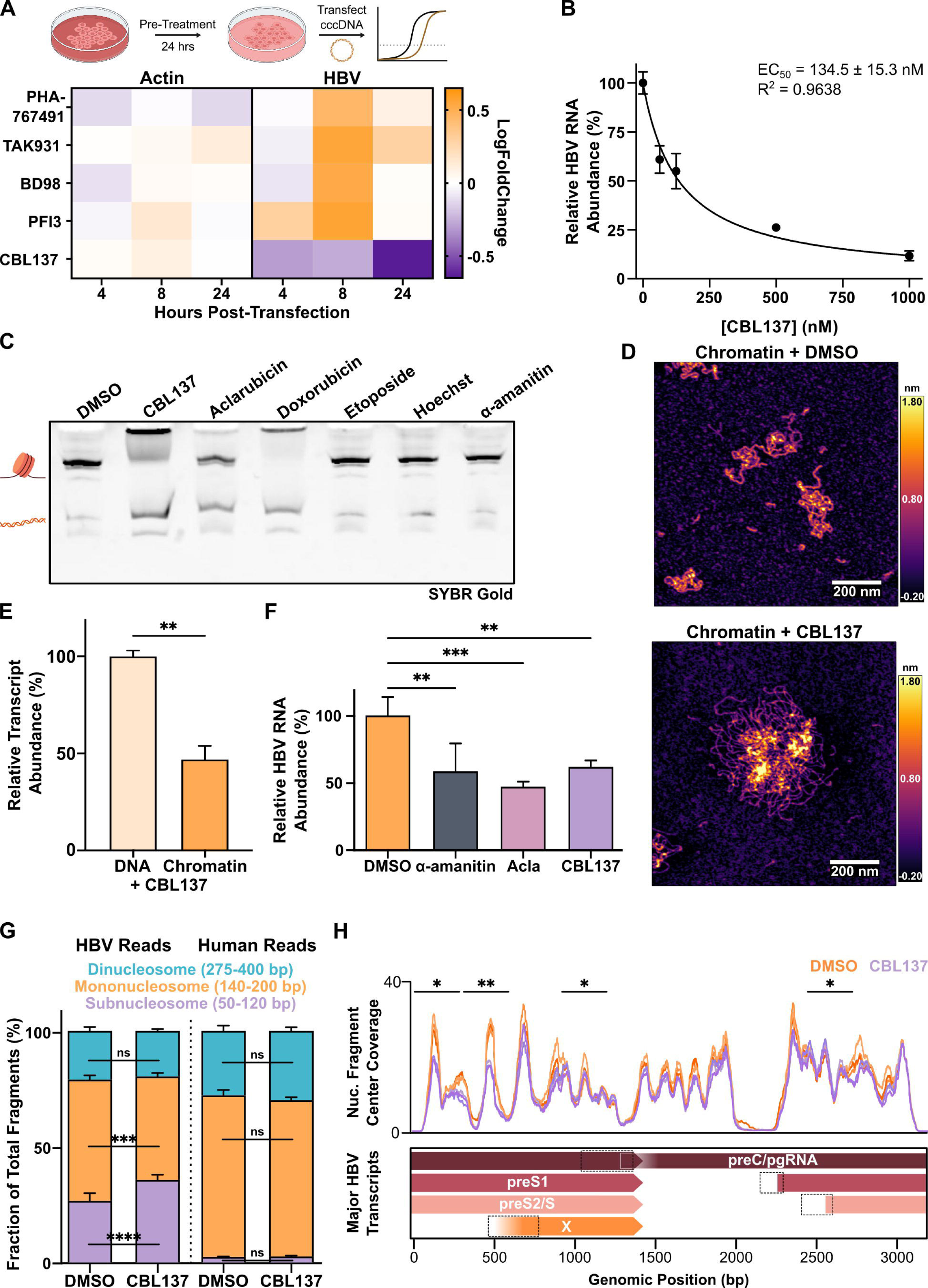
Nucleosome destabilizing drugs inhibit viral transcription in cells and disrupt cccDNA integrity *in vitro*. (A) Heatmap depicting log-fold change of HBV or β-actin mRNA levels, normalized against GAPDH, in cccDNA-transfected HEK 293T cells treated for 24 hours with the indicated molecules (1µM PHA-767491, 300 nM TAK-931, 2.5 µM BD98, 1 µM PFI-3, 500 nM CBL137). Data represent 2 biological replicates. (B) Dose-dependence of HBV RNA levels in HEK293T cells 24-hours after cccDNA transfection and concurrent treatment with the indicated concentration of CBL137. (C) Native PAGE gel stained with SYBR Gold to assess the stability of recombinant mononucleosomes following incubation with the indicated molecules (all at 10 µM concentrations). (D) Representative atomic force micrographs of recombinant HBV dslDNA chromatin arrays treated with CBL137 or a vehicle control prior to AFM imaging. (E) Quantification of X transcript produced following *in vitro* transcription of empty or chromatinized chimeric X-601 template DNA using HepG2 nuclear extract, relative to an internal control from an unchromatinized template under control of the CMV promoter, in reactions where 1 µM CBL137 was added. (F) Quantification of relative HBV RNA 24 hours after transfecting HEK293T cells with recombinant cccDNA and concurrent treatment with DMSO (vehicle), 500 nM α-amanitin (positive control), 125 nM CBL137, or 50 nM aclarubicin. (G) Quantification of the fraction of total subnucleosome-(50-120 bp), mononucleosome-(140-200 bp), or dinucleosome-sized (275-400 bp) DNA fragments mapping to either the HBV or human genomes following MNase-seq of HepAD38 nuclei after 48 hour treatment with 500 nM CBL137 or a vehicle control. (H) Coverage of centers of mononucleosome-length DNA fragments mapping to the HBV genome from G, where indicated 300 bp regions show a significant change in fragment counts between mock- and CBL137-treated samples. Data represent 2 (A-D), 3 (E, G-H), or 4 (F) independent experiments/biological replicates as means ± SD and were analyzed by 1-way ANOVA with Dunnett’s multiple comparison test (F), Welch’s t-test (E), and 2-way ANOVA with Tukey’s multiple comparison test (G), and two-tailed two-sample t-test (H). *P < 0.05, **P < 0.01, ***P < 0.001.

Motivated by these promising results, we sought to better understand how CBL137 inhibits HBV transcription and the role of chromatin in this process. CBL137 was first described as an inhibitor of the histone H2A-H2B chaperone FACT complex, capable of redistributing it in nuclei to activate p53 and suppress NF-kB.^46^ More recently its mechanism of FACT inhibition has been elucidated, whereby CBL137 intercalates into nucleosomal DNA, disrupting nucleosome integrity and trapping FACT on destabilized chromatin, preventing it from chaperoning H2A-H2B dimers elsewhere across the genome (e.g., at transcription sites).^50^ This led us to test whether the anti-HBV effect of CBL137 is due to chromatin destabilization rather than formal inhibition of FACT. We first tested the ability of CBL137 and other molecules previously shown to induce histone eviction from chromatin (the anthracycline topoisomerase II inhibitors aclarubicin and doxorubicin) to destabilize mononucleosomes *in vitro.*^51^ Reconstituted 601 mononucleosomes incubated with either CBL137, doxorubicin, and aclarubicin treatment exhibited increased free DNA and/or aggregated chromatin in gel wells when samples were resolved by native gel electrophoresis, whereas the chemically distinct topoisomerase II inhibitor etoposide, the DNA-binding dye Hoechst, and the RNAPII inhibitor α-amanitin did not (Fig. 4C). To characterize nucleosome destabilization more quantitatively, we implemented a thermal stability assay to measure the binding of a hydrophobic fluorescent dye as H2A-H2B dimers and (H3-H4)2 tetramers are evicted from nucleosomal DNA with increasing temperatures. Concordant with our native gel results, we found that CBL137, doxorubicin, and to a lesser extent aclarubicin, significantly decreased the melting temperature for H2A-H2B dissociation (Tm1), with CBL137 also decreasing (H3-H4)2 melting temperature (Tm2) (Fig. S4B). All three mononucleosome destabilizing molecules were similarly able to induce a marked destabilization of reconstituted HBV chromatin arrays, causing chromatin fibers to resolve as smears rather than distinct bands on a native gel (Fig. S4C). We subsequently visualized the aggregative effects of CBL137 on HBV chromatin using AFM (Fig. 4D). We likewise measured decreased thermal stability in reconstituted Lin601^18^, Cir601^18^, and X-601 chromatin arrays treated with CBL137, confirming the molecule’s ability to destabilize various chromatin fibers (Fig. S4D). Importantly, we found that the addition of CBL137 to *in vitro* transcription reactions with DNA or chromatinized X-601 template resulted in significantly reduced transcription from chromatinized templates, using both HeLa and HepG2-derived extracts (Fig. 4E, Fig. S4E).

Based on these results, we next tested if treating cells with anthracyclines could phenocopy the effects of CBL137 treatment on viral transcription, focusing on aclarubicin rather than doxorubicin to avoid potential confounding effects from doxorubicin-induced DNA damage.^51^ Indeed, aclarubicin treatment induced a significant decrease in total HBV transcription 24 hours after cccDNA transfection into HEK293T cells and concurrent drug treatment. (Fig. 4F). Interestingly, previous reports have implicated anthracyclines as having potential anti-HBV effects through the inhibition of cccDNA synthesis, but these results suggest they may have additional mechanisms of action.^52^ Curious about the genomic distribution of destabilized chromatin within cccDNA, we performed additional MNase-seq experiments on HepAD38 cells treated with CBL137 or a vehicle control. We detected decrease in the proportion of mononucleosomes-sized fragments and corresponding increase in the proportion of sub-nucleosome-sized fragments (50-140 bp) mapping to the HBV genome in cells treated with CBL137, but no such change for DNA fragments mapping to the human genome (Fig. 4G, S4F). Examining whether disruption of chromatin integrity was evenly distributed across the HBV genome, we found that although most loci exhibited at least a slight decrease in nucleosome-sized DNA fragments, parts of both the X promoter (nt 301-600) and gene body (901-1200) exhibited a significant decrease in nucleosome-sized DNA fragments (Fig. 4H). For an orthogonal validation, we transfected HepG2 cells with recombinant cccDNA, with or without concurrent CBL137 treatment for 24 hours, and subjected them to ChIP-seq, probing for the distribution of H3K4me3 and RNAPII, as well as the core histones H2A, H2B, and H3 along cccDNA. Human-aligned reads from all samples appeared as expected, with TSS-associated peaks for H3K4me3 and RNAPII but relatively linear distributions for core histones (Fig. S4G). Sequencing reads for all targets mapped along the entire viral genome, and quantitative analysis of normalized read count from mock- or CBL137-treated samples revealed a marked redistribution of H3K4me3, RNAPII, and core histone enrichment at specific genomic loci (Fig. S4H). In particular, we observed a significant decrease in RNAPII, H2A, and H2B reads in part of the genomic region containing the X promoter and TSS following CBL137 treatment (Fig. S4I). Collectively, these results indicate that destabilization of cccDNA chromatin integrity, particularly around the X promoter region, can impede HBV transcription.

### CBL137 reduces viral transcription, antigen secretion, and genome replication in models of infection

Encouraged by the possibility that CBL137 may pose a potential therapeutic opportunity based on its reduction of HBV transcription, we next tested this in additional cellular models of infection. We first performed similar transfection experiments as before to introduce recombinant cccDNA into HepG2 cells and measured the impact of CBL137 treatment on overall HBV transcription. As expected, total viral RNA levels were reduced by over 50 % in CBL137-treated cells at 4 hours post-transfection, and almost completely abolished by 24 hours relative to GAPDH (Fig. 5A). Transcript-specific RT-qPCR analysis from these samples revealed that at both time points, in addition to a global decrease in viral RNA abundance, CBL137 treatment also significantly reduced the relative proportion of X transcript (Fig. 5B). We next tested if long-term CBL137 treatment could impact viral transcription in HepAD38 and HepG2.2.15 cells. We first identified a milder CBL137 dose regime compared to those used in experiments with early endpoints that did not inhibit cell viability with prolonged treatment (Fig. S5A, Table S1). Both cell lines were then cultured in media containing 50 nM CBL137 for nearly two weeks, harvesting samples periodically for analysis. We found that under these conditions, total HBV RNA was reduced within 6 days and for up to 11 days without significantly harming cell viability (Fig. S5B). We noted that the magnitude of the reduction was less than we observed previously, potentially due to differences in the transcriptional regulation of the integrated genome copies in these cell lines, which rely on strong promoters to drive expression.

**Figure 5.**
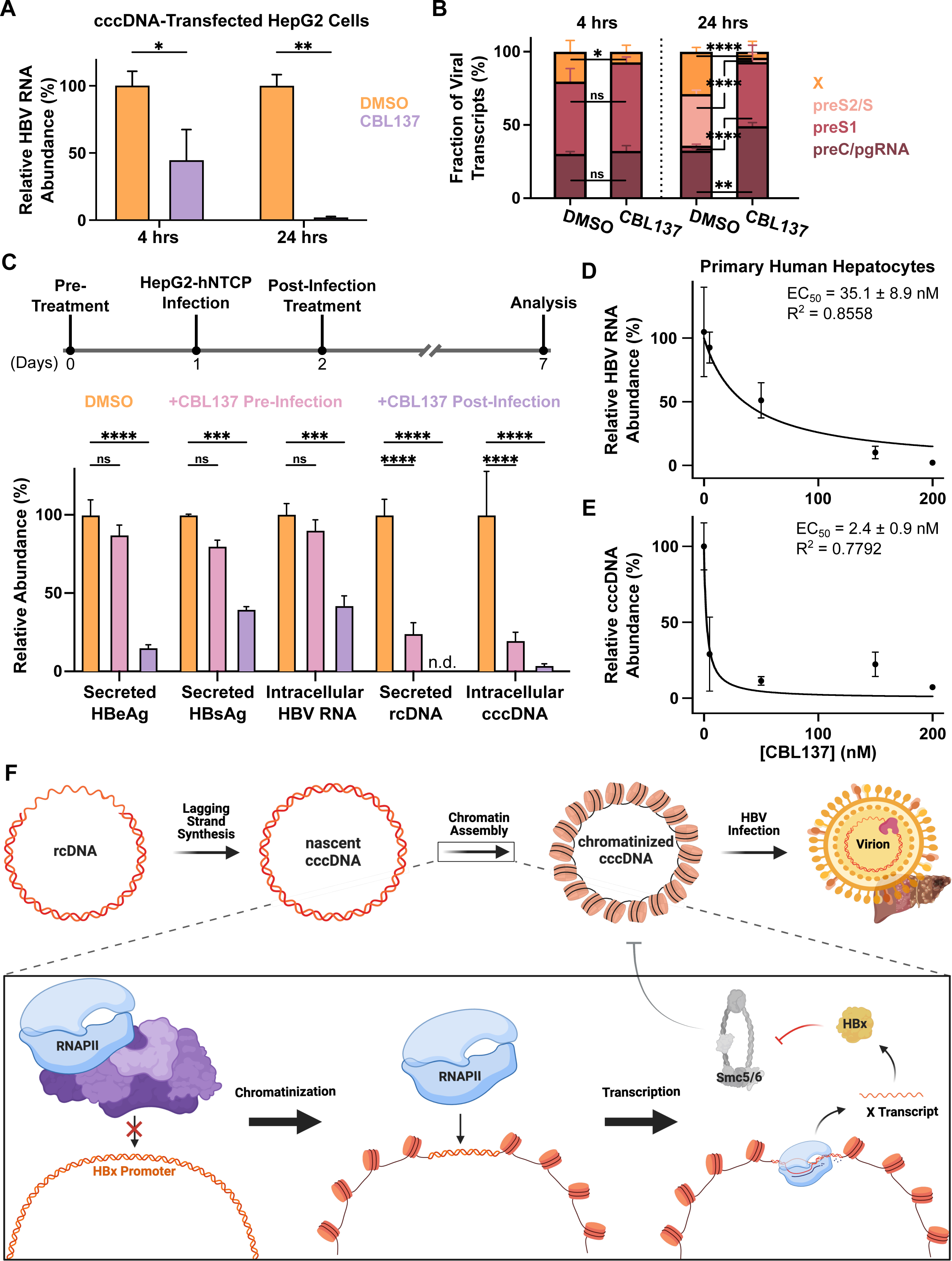
CBL137 inhibits viral transcription, antigen secretion, and genome replication in hepatocyte models of HBV infection. (A) Quantification of relative total HBV RNA from HepG2 cells 4- or 24-hours after transfection with recombinant cccDNA and concurrent treatment with DMSO or 500 nM CBL137. (B) Quantification of the proportion of total HBV RNA comprised by each of the four major viral transcripts from samples in A. (C) Quantification of secreted HBV surface antigen (HBsAg), secreted antigen (HBeAg), intracellular total RNA, secreted rcDNA, or intracellular cccDNA from HepG2-NTCP cells following infection and either 24 hour pre-infection treatment or 5 day post-infection treatment with 125 nM CBL137. (D) Dose-dependence of total HBV RNA abundance in primary human hepatocytes infected with HBV for 5 days and subsequently treated with the indicated concentrations of CBL137 for 2 days. (E) Dose-dependence of intracellular cccDNA abundance in primary hepatocytes cultured and treated as in D. (F) Proposed model illustrating recognition of the chromatinized X gene promoter as a driving force behind transcription of the X gene during the genesis of infection. All data are means ± SD of 3 biological replicates and were analyzed by Welch’s t-test (A) and 2-way ANOVA with Tukey’s (B) or Dunnett’s (C) multiple comparisons tests. *P < 0.05, **P < 0.01, ***P < 0.001, ****P <0.0001.

We next turned to HepG2-NTCP cells to test the effects of CBL137 on *bona fide* infections with virus collected from the supernatant of HepG2.2.15 cells. Cells were either pre-treated with 125 nM CBL137 starting one day prior to infection or one day after infection, and media was collected from each sample after 6 days for analysis of viral antigen production. Post-infection treatment with CBL137 led to significant reduction in viral antigen secretion and intracellular viral RNA abundance, whereas the pre-treatment only yielded negligible changes compared to mock-treated cells (Fig. 5C). Dual pre- and post-infection treatment had no significant differences from post-infection treatment only (Fig. S5C). We also investigated whether these treatment regimens could reduce viral genome replication in this model system. Notably, even one-day pre-treatment of cells with the drug was able to significantly diminish both secreted rcDNA titer in cell media and intracellular cccDNA levels to approximately 25 % of those in untreated cells. Even more strikingly, post-infection treatment for 5 days led to near-total elimination of secreted rcDNA from media and reduced cccDNA levels to only 5 % that of untreated cells (Fig. 5C). Together, these results suggest that treatment with CBL137 can not only impede viral transcription during early stages of infection, but also reduce viral protein secretion and genome replication. We speculate that the observed dramatic decrease in total cccDNA copy number despite a comparably smaller decrease in the total HBV RNA or secreted antigen abundance in CBL137-treated cells is due to diminished genome recycling and *de novo* secondary infection events, which have been shown to be important for the maintenance of cccDNA levels.^53^

Finally, we assessed the efficacy of CBL137 at inhibiting HBV in primary human hepatocytes (PHHs), the gold standard *in vitro* model for HBV replication and infection. PHHs were cultured, infected with HBV for 5 days, and then treated with CBL137 for 48 hours before analysis. In agreement with our results from infected HepG2-NTCP cells, CBL137 treatment induced a dose-dependent reduction of secreted antigen and viral nucleic acid abundance with doses at or below 200 nM (Fig. 5D-E, S5D). Importantly, CBL137 treatment did not detectably harm primary hepatocyte viability or function, as measured by alpha-1-antitrypsin and albumin secretion, in this dose regime (Fig. S5E-G). These results demonstrate CBL137 as an effective inhibitor of HBV transcription and replication that may pose a potential new therapeutic avenue to treat HBV infection.

## Discussion

Hepatitis B Virus is an ongoing global health problem responsible for nearly one million deaths per year. Despite decades of study, a significant knowledge gap remains with respect to how the chromatinization status of the virus’s cccDNA minichromosome relates to its transcriptional capacity. Here, we report a robust platform to generate reconstituted HBV minichromosomes that has laid the foundation for mechanistic investigations of a critical chromatin species in HBV pathogenesis. This recombinant, functional cccDNA serves as a powerful tool to perform high resolution biochemical investigation of the minichromosome as well as to surmount the temporal heterogeneity and abundance limitations of viral infection, enabling closer study of early infection time points. We used this system to perform mechanistic investigations that led to a model wherein chromatin occupancy on cccDNA, particularly in the X gene promoter and ORF, plays a pivotal role in its early transcription and thus HBV infection (Fig. 5F). We corroborated our *in vitro* results with a suite of transcriptomic, nucleosome mapping, and pharmacological experiments, and have shown that this phenomenon may have important clinical implications as a therapeutic treatment for chronic HBV infection in the future.

The results of this work also contribute insights into the basic biology of HBV infection. The rapid X transcription that we observed by RNA-seq (Fig. 3D, E) is consistent with prior studies of HBV transcript kinetics, as well as the intuitive notion that for infection to progress HBx must be generated rapidly to induce the degradation of Smc5/6 that would otherwise suppress infection.^13,14,16^ Several reports have found X RNAs in viral particles circulating in patient sera samples and in the supernatant of infected cell lines, leading to an outstanding question in the field - whether X mRNA is delivered to naïve hepatocytes alongside rcDNA.^15,17,54^ Contrary to this idea, our data show that the introduction of intact cccDNA to wild-type cells is sufficient for rapid X transcription, suggesting that the packaging of X RNA into viral capsids may be non-specific, as has been observed for host transcripts in HBV and other viral infections.^55,56^ The specific details of how X can be so rapidly and specifically expressed prior to Smc5/6-induced silencing remains a topic for future studies with this system.

Our work reveals a new aspect of the transcriptional regulation of the X gene. The DNA sequence of the X promoter is distinguished by its lack of typical well-annotated core promoter elements (e.g., TATA box, Initiator [Inr], or downstream promoter element [DPE] sequences). Instead, it includes two core promoter elements, XCPE1 and XCPE2, that drive transcription initiation and inspired the identification of similar elements at a subset of TATA-less promoters across the human genome.^57,58^ On the contrary, a canonical TATA box is present in the preS1 promoter, and TATA-like motifs coupled with Inrs are in the preC and pgRNA promoters.^59,60^ While the preS2 promoter lacks a TATA box, its promoter resembles the SV40 late promoter and has been well-characterized.^60^ Luciferase-based reporter assays have previously shown that the X gene promoter is comparatively weaker than those of other HBV transcripts, a finding also supported by the weaker signal for the X transcript commonly observed in Northern blot assays of infected cells.^19,61^ The exact mechanism by which the presence of nucleosomes enhances the X gene promoter’s otherwise weak activity remains to be determined, but one possible explanation to reconcile these findings may arise from studies on the PIC-Mediator complex, a mega-complex responsible for eukaryotic transcription initiation. Earlier biochemical work revealed that PIC-Mediator can facilitate transcription through a chromatinized promoter.^62^ Recent structural studies have further found that PIC-Mediator can directly bind to the +1 nucleosome, and in doing so specifically enhance transcription from TATA-like and TATA-less promoters.^38^ Thus, a hypothesis arising from these combined observations, to be tested in future studies, is that nucleosome assembly at the X TSS may drive transcription by allowing PIC-Mediator to recognize a promoter that otherwise cannot efficiently recruit the basal transcription machinery based on DNA sequence alone.

Our results indicate that pharmacological perturbation of chromatin integrity obstructs HBV infection, providing a new potential therapeutic avenue while also underscoring the need to characterize the epigenomic landscape of cccDNA more robustly. Although such efforts have been undertaken in duck HBV,^27^ we describe some of the first high-resolution characterization of nucleosome positioning across the HBV genome in mammalian cells (Fig. 3A). Similarly sparse is our knowledge of histone modifications and variants in cccDNA. Other studies have found that cccDNA is decorated overwhelmingly with active transcription-associated PTMs, and mapped positions of H3K4me3, H3K27ac, and H3K122ac, among others, along the genome.^19,63^ Lower resolution methods have also been applied to identify other PTMs, such as H4ac, and more recently the histone variant H3.3 within cccDNA.^18,64–66^ Additional studies have reported that transcriptionally active cccDNA localizes to the more accessible “A-compartment” of chromatin in infected cells,^67,68^ possibly explaining its relatively high susceptibility to MNase digestion even without CBL137 treatment. As more is known about the chromatin of cccDNA, therapeutic opportunities may become available to target the minichromosome if not for degradation, then to silence it and establish a functional cure.^69^

Recent drug discovery and development efforts have focused on generating novel chemical tools and finding alternate targets besides the HBV polymerase, the current primary target for antiviral therapies. For example, core protein allosteric modulators (CpAMs) have emerged as encouraging potential drugs to disrupt capsid assembly and cccDNA maintenance.^70,71^ Even newer drugs have emerged in recent years such as RG7834, which inhibits RNA binding proteins that stabilize HBV transcripts,^72–74^ or the recently reported cccDNA inhibitor ccc_R08, for which a mechanism of action has not yet been determined.^75^ We believe that our results showing the potent inhibitory effects of CBL137 on HBV replication expand the list of potential avenues towards new HBV therapeutics. Finally, the chromatin destabilizing activity of CBL137 merits further investigation to test if it could prove similarly useful to target or study other chromatinized DNA viruses including herpesviruses, papillomaviruses, and adenoviruses. Indeed, recent studies have reported encouraging results for the use of CBL137 as a potential latency-reversing agent in HIV-1 infection, and even as a lead for drug discovery efforts against human African trypanosomiasis, further underscoring the potential of CBL137 as a therapeutic against various infectious diseases.^76,77^

### Limitations of the Study

In this study, we reconstituted the HBV cccDNA minichromosome for biophysical and biochemical studies, which revealed a chromatinization-dependent increase in the transcription of the gene encoding a key viral protein, HBx. We then proceeded to find that the chromatin destabilizing small molecule CBL137 inhibits not only viral transcription *in vitro*, but also antigen secretion and genome replication in cellular models of HBV infection. While we propose that the mechanism of CBL137’s anti-HBV activity is due to the disruption of chromatin architecture at the X promoter, e.g. by possibly disrupting the -1/+1 nucleosome to prevent binding of the PIC-Mediator supercomplex, extensive additional epigenomic and transcriptomic studies will be needed to more definitively test this and competing hypotheses. Additionally, we cannot rule out whether disruption of the FACT complex by CBL137, as it was first described to do, does not contribute to its activity. Technical limitations arising from the FACT complex’s essentiality in proliferating cell lines prevented us from carrying out knockdown/knockout experiments to test whether CBL137 exhibits the same activity in the absence of FACT.^78^ Likewise, CBL137-mediated disruption of spatial genome organization (e.g., looping by CTCF or torsional strain) are also potential mechanisms of inhibition that remain to be more thoroughly addressed.^66,79–81^

## Acknowledgments

The authors thank members of the David, Schwartz, and Risca labs for their support. We thank B. Y. Winer for helpful discussions and feedback, as well as B. Wang and K. Manova-Todorova for technical assistance. We acknowledge the use of the MSKCC Integrated Genomics Operation, Molecular Cytology Core Facility, and Center for Epigenetics Research, supported by the NCI Cancer Center Support Grant (CCSG, P30CA08748), Cycle for Survival, and the Marie-Josée and Henry R. Kravis Center for Molecular Oncology; and of the Genomics Resource Center and High Performance Computing Cluster at Rockefeller University. The Epigenetics Research Innovation Lab is partially funded by the Metropoulos Family Foundation and the Gladys & Richard Harriman Foundation. This research was supported by the National Institutes of Health (grants T32GM115327-Tan and F99CA264420 to N.A.P.; F32GM140551 to A.M.; T32GM136640-Tan to A.A.L.; DP2GM150021 to V.I.R.; R01AA027327, R01AI107301, and R01DK121072 to R.E.S), the National Science Foundation (graduate research fellowship 2017239554 to N.A.P.), the Josie Robertson Foundation (to Y.D), Alfred Sloan Research Foundation (Y.D.) and the MSKCC Center for Epigenetics Research (to Y.D.), the Rita Allen Foundation (scholar award to V.I.R.), the Irma T. Hirschl Trust (Monique Weill-Caulier career scientist award to V.I.R. and research scholar award to R.E.S.), the Starr Cancer Consortium (I16-0058 to V.I.R.), the V Foundation for Cancer Research (V Scholar award to V.I.R.), the Stavros Niarchos Foundation (SNF) as part of its grant to the SNF Institute for Global Infectious Disease Research at The Rockefeller University (grant to A.R.M., sponsored by V.I.R.), the American Cancer Society (fellowship PF-23-1034949-01-CCB to J.R.), the United States Department of Defense (W81XWH-21-1-0978 to R.E.S.), and the Paul G. Allen Family Foundation (UWSC13448 to R.E.S.).

## Author contributions

Conceptualization, N.A.P., R.E.S, and Y.D.; Methodology, N.A.P., A.M., Y.B., T.B., S.C.F., and V.I.R.; Investigation, N.A.P., A.M., Y.B., T.B., J.R., S.C.F., A.A.L., C.L., P.-J.H., R.K., and V.I.R.; Visualization, N.A.P., A.M., and J.R.; Writing – Original Draft, N.A.P. and Y.D.; Writing – Review & Editing, N.A.P., A.M., Y.B., T.B., J.R., S.C.F., A.A.L., V.I.R., R.E.S., and Y.D.; Supervision, V.I.R., R.E.S., and Y.D.; Project Administration, N.A.P., V.I.R., R.E.S., and Y.D.; Funding acquisition: N.A.P., A.M., V.I.R., R.E.S., and Y.D.

## Supplemental Figure Titles and Legends

**Figure S1.**
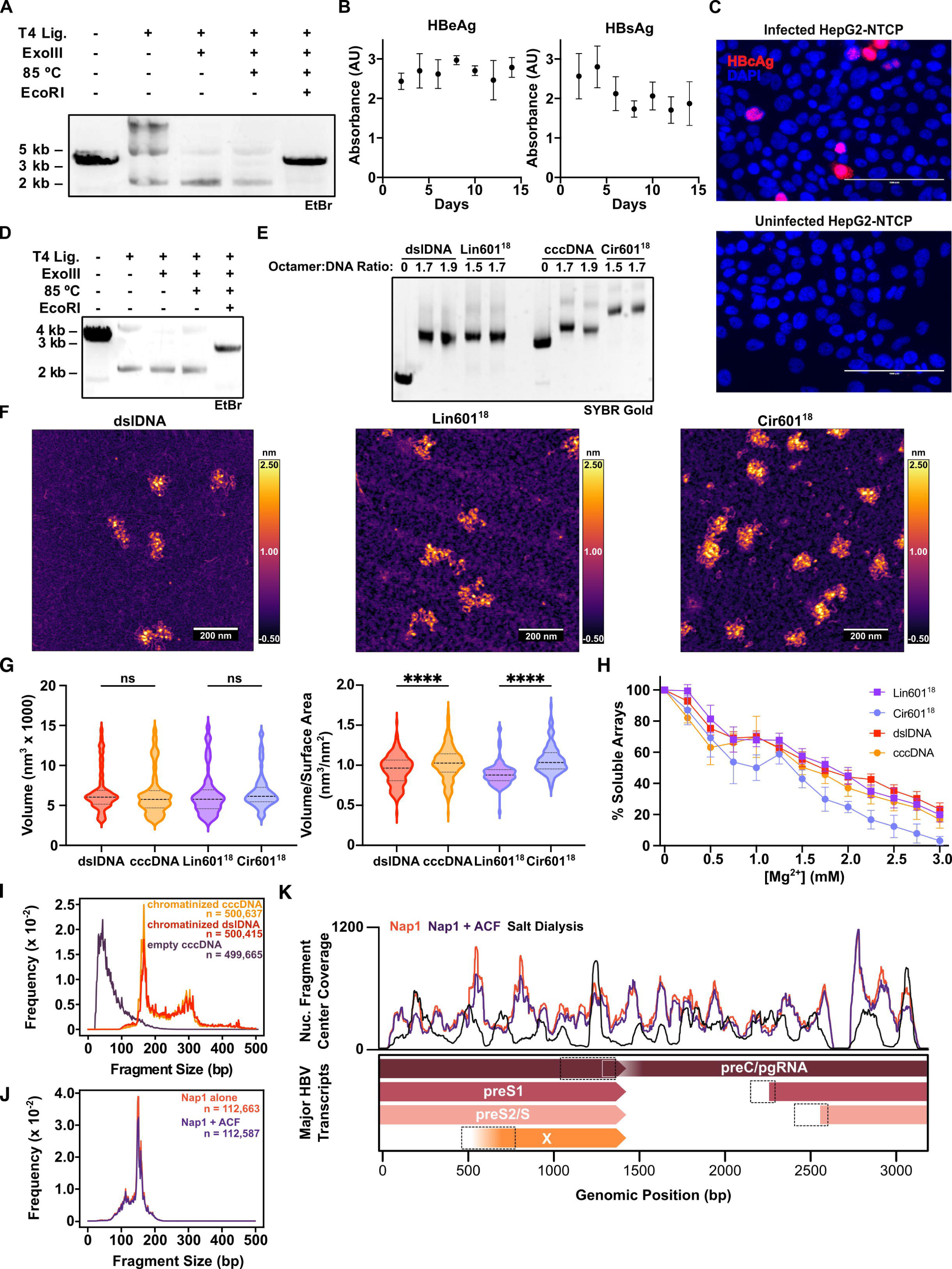
Preparation, validation, and comparison of recombinant HBV and 601^18^ chromatin fibers and minicircles, related to Figure 1. (A) Agarose electrophoresis gel stained with ethidium bromide to illustrate conversion of dslDNA (lane 1, 3.2 kb band) into cccDNA (2.1 kb band) following ligation (lane 2), exonuclease digestion of linear intermolecular ligation products (lane 3), and incubation at 85 °C for 5 minutes to melt nicked or unligated species (lane 4). Subsequent digestion with EcoRI reverts cccDNA back into dslDNA (lane 5). (B) ELISA of HBV secreted (HBeAg) and surface (HBsAg) antigens following transfection of HepG2 cells with recombinant cccDNA. (C) Immunofluorescence staining of HBV capsid protein (HBcAg) on mock- or HBV-infected HepG2-NTCP cells. (D) Agarose electrophoresis gel stained with ethidium bromide to illustrate conversion of Lin601^18^ (lane 1, 3.2 kb band) into plasmid to Cir601^18^ (2.1 kb band) following ligation (lane 2), exonuclease digestion (lane 3), and incubation at 85 °C for 5 minutes (lane 4). Subsequent digestion with EcoRI reverts the minicircles back into Lin601^18^ (lane 5). (E) EMSA of linear and circular HBV and 601 DNA and chromatinized samples on agarose-polyacrylamide native gel. (F) Representative AFM micrographs of dslDNA, Lin601^18^, and Cir601^18^ chromatin fibers. (G) Volume (left) and volume-surface area ratio (right) quantifications of the four chromatin species prepared. (H) Magnesium-dependent self-association assay with all four chromatin species, illustrating similar precipitation/aggregation behaviors. (I) Fragment length distributions of MNase-digested salt-gradient dialysis-assembled cccDNA (orange) or dslDNA (red) chromatin arrays, or empty cccDNA (navy). (J) Fragment length distributions following MNase digest of dslDNA chromatin fibers assembled by the histone chaperone Nap1 (orange) or Nap1 with the chromatin remodeler ACF (navy). (K) Coverage of centers of mononucleosome-length (140-200 bp) DNA fragments from J overlaid against MNase-seq coverage of dslDNA fibers assembled by salt-gradient dialysis (black) above a schematic of the four major HBV transcripts. Dotted boxes indicate promoter regions for each transcript, in genomic coordinate order: enhancer I/X promoter, enhancer II/basal core promoter, S1 promoter, S2 promoter.

**Figure S2.**
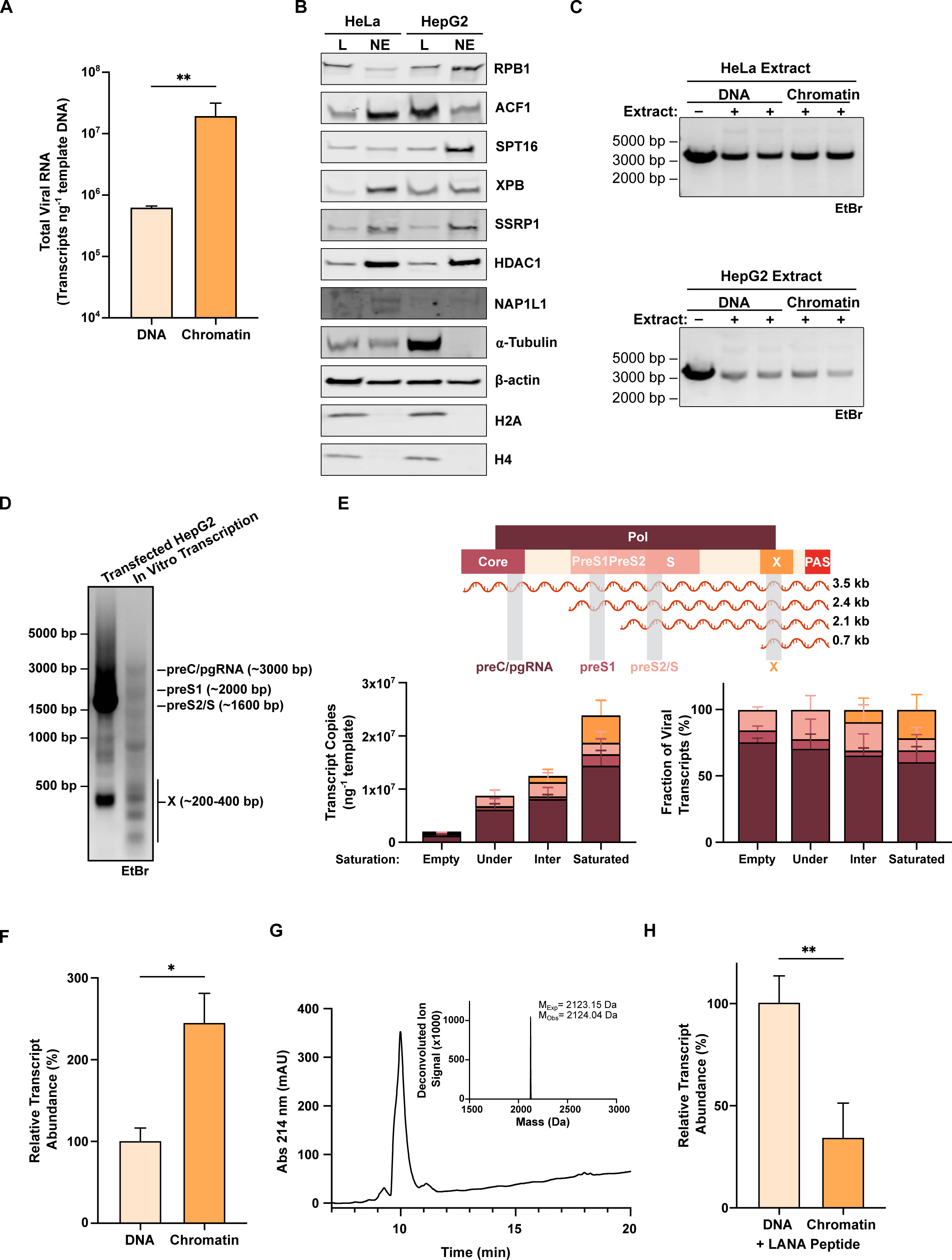
Characterization and additional analysis following *in vitro* transcription of HBV chromatin, related to Figure 2. (A) Absolute quantification of total HBV RNA following *in vitro* transcription (IVT) of empty or chromatinized cccDNA in commercial HeLa cell nuclear extract. (B) Immunoblots probing the presence of the indicated targets in whole cell lysates (L) or nuclear extracts (NE) prepared from HeLa or HepG2 cells. (C) Agarose electrophoresis gel stained with ethidium bromide to assess the integrity of empty or chromatinized Lin601^18^ DNA incubated in HeLa (top) or HepG2 (bottom) nuclear extracts as for IVT reactions. (D) Agarose electrophoresis gel stained with ethidium bromide to visualize HBV transcripts following full-length, 5’RACE of RNA isolated from HepG2 cells 24 hours after transfection with cccDNA (left) or from IVT reactions of chromatinized cccDNA in HepG2 nuclear extract (right). Note all labeled transcripts appear approximately 500 bp shorter due to the positioning of gene-specific primers used for 5’RACE, as described previously.^17^ (E) Schematic depiction of the four major viral transcripts and primer pairs used to quantify overlapping and non-overlapping regions therein for transcript-specific RT-qPCR analysis (top), and absolute quantification (bottom left) and relative proportion of total viral RNA (bottom right) of HBV transcripts following IVT of empty, undersaturated, intermediately, or fully saturated cccDNA templates in HeLa nuclear extract. (F) Quantification of X transcript produced following IVT of empty or chromatinized chimeric X-601 template DNA, relative to an internal control from an unchromatinized template under control of the CMV promoter, in HeLa nuclear extract. (G) Analytical LC-MS chromatogram and deconvoluted mass spectrum (inset) to characterize synthetic LANA peptide. (H) Quantification of X transcript produced as in F, with 1 µM LANA peptide added to all reactions. Data are representative of 2 (B-D) and 3 (A, E, F, H) replicates ± SD and were analyzed by Welch’s t-test (A, F, H). *P < 0.05, **P < 0.01, ***P < 0.001, ****P <0.0001.

**Figure S3.**
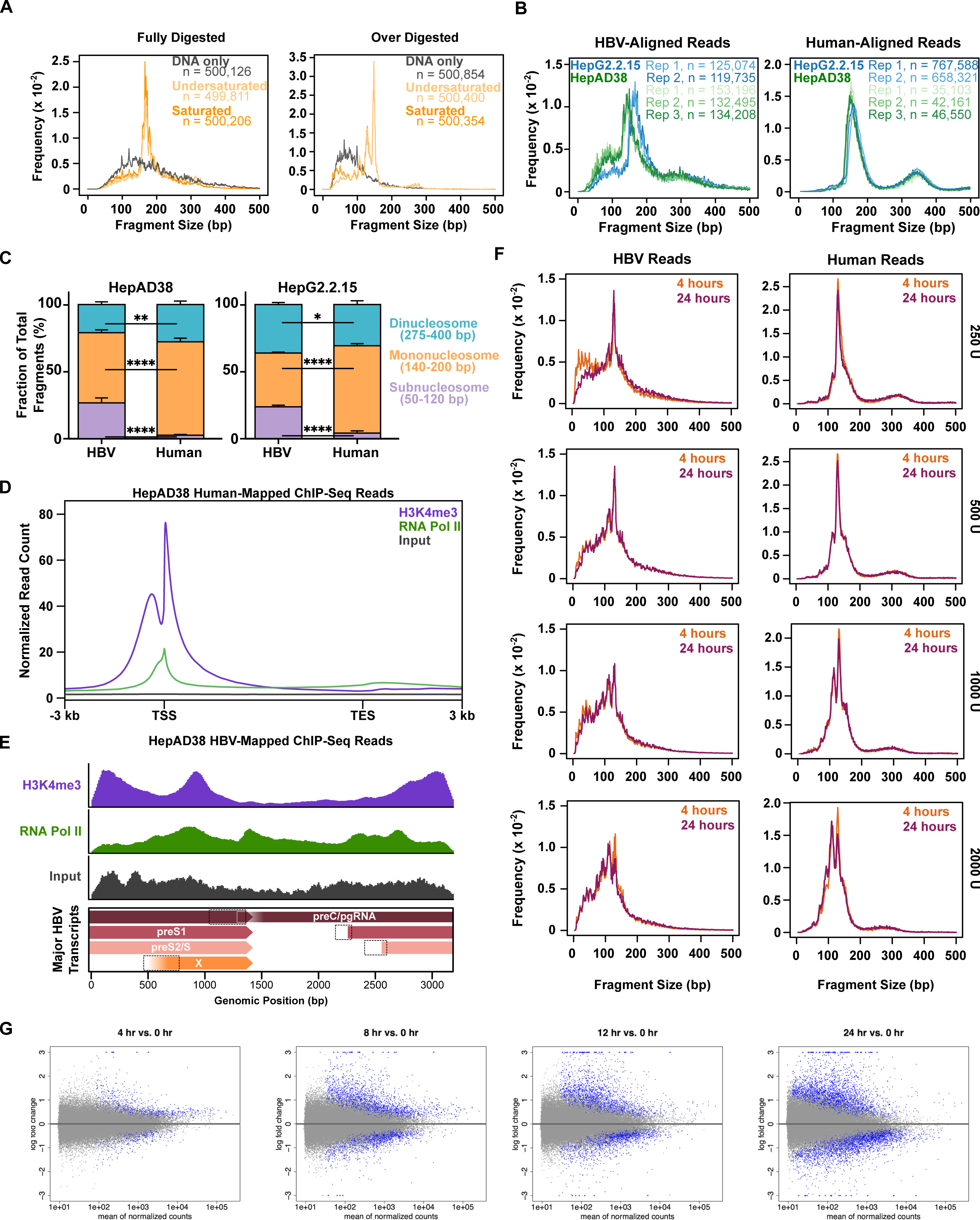
MNase-seq fragment-length distributions, ChIP-seq analysis, and human RNA-seq analysis, related to Figure 3. (A) Fragment length distributions following complete MNase digest (left) or overdigestion (right) of differentially saturated cccDNA chromatin arrays depicted in Figure 3A. (B) Fragment length distributions of HBV- and human-mapped reads following MNase digest of HepG2.2.15 and HepAD38 nuclei. (C) Quantification of the fraction of total DNA fragments mapping to either the HBV or human genomes from B that fall into subnucleosome-(50-120 bp), mononucleosomes-(140-200 bp), or dinucleosome-length (275-400 bp) bins.) (D) Metagene profile composite plots for H3K4me3 and RNAPII along the human genome in HepAD38 cells, with signal normalized to 10 million uniquely mapped reads across the entire genome. (E) Read density profiles of H3K4me3 and RNAPII distributions along the HBV genome in HepAD38 cells, all normalized to 10 million uniquely mapped reads and scaled to 1 million reads on the y-axis. (F) Fragment length distributions of HBV- and human-aligned MNase-seq fragments from HEK293T cells transfected with cccDNA and harvested after 4 or 24 hours using different quantities of MNase for sample preparation. (G) Pairwise analysis of human gene expression changes in non-transfected HEK293T cells and cccDNA-transfected cells revealed few differentially expressed genes (defined as log fold change > 2and p <0.05) at 4 hrs (0 down, 1 up), 8 hrs (2 down, 6 up), 12 hrs (9 down, 18 up), and 24 hrs (36 down, 49 up) following transfection. Data are representative of 2 (A, D-F) or 3 (B-C, G) replicates.

**Figure S4.**
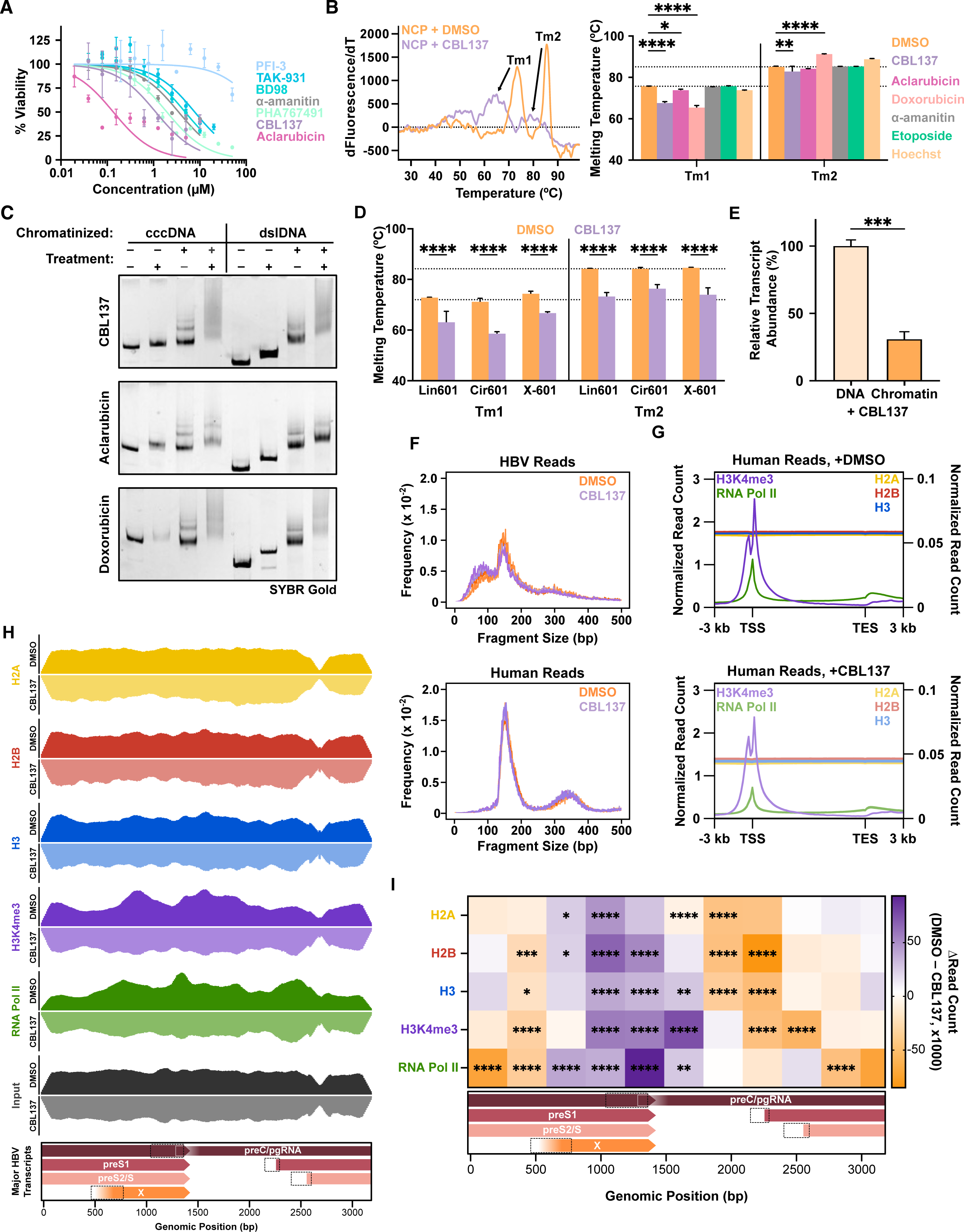
Additional validation of chromatin destabilization *in vitro* and in cells, related to Figure 4. (A) HEK293T cells were treated with the indicated drug concentrations for 48 hours before harvesting and measuring cell viability with CellTiter-Glo assay. (B) Representative thermal stability curve depicting the first derivative of sample fluorescence with respect to temperature for DMSO-(orange) or 10 µM CBL137-treated (violet) 601 mononucleosomes (left) and quantification of mononucleosome melting temperatures following incubation with the indicated molecules (right). Canonical melting peaks for H2A/H2B dimer dissociation (Tm1) and (H3/H4)2 tetramer dissociation (Tm2) from DNA are indicated for DMSO sample and arrows depict the shift of these peaks in CBL137 treated samples. (C) Representative agarose-polyacrylamide native EMSAs of intact recombinant cccDNA and dslDNA chromatin fibers following 1-hour incubation with a 10 µM dose of the following drugs: CBL137 (top), aclarubicin(middle), and doxorubicin (bottom). Representative of 2 independent experiments. (D) Quantification of chromatin array melting temperatures (Tm1, left and Tm2, right) for the indicated species following incubation with DMSO or 10 µM CBL137. (E) Quantification of X transcript produced following *in vitro* transcription of empty or chromatinized chimeric X-601 template DNA using HeLa nuclear extract, relative to an internal control, in reactions where 1 µM CBL137 was added. (F) Fragment length distributions for HBV-(top) and human-aligned (bottom) MNase-seq reads from data depicted in Figure 4G-H. (G) Metagene profile composite plots for H2A, H2B, H3, H3K4me3, and RNAPII along the human genome in HepG2 cells 24 hours after transfection with recombinant cccDNA and concurrent treatment with DMSO or 500 nM CBL137, with signal normalized to 10 million uniquely mapped reads across the entire genome. Left y-axis reflects normalized read count for H3K4me3 and RNAPII, right y-axis reflects that of the core histones. (H) Read density profiles of HBV-aligned ChIP-seq data for the indicated antibodies in DMSO-(top, solid) or CBL137-treated (bottom, faded, mirrored across the x-axis) samples, all normalized to 10 million uniquely mapped reads and scaled to 600,000 reads on the y-axis. (I) Heatmap depicting changes in normalized ChIP-seq read count of 300 bp regions of the HBV genome between mock- and CBL137-treated samples, where stronger purple color indicates more reads in DMSO-treated samples. Data represent means of 2 (G-I) or 3 (A-F) replicates ± SD and were analyzed by 2-way ANOVA with Dunnett’s multiple comparison test (B) or Tukey’s multiple comparisons test (D, I), and Welch’s t test (E). *P < 0.05, **P < 0.01, ***P < 0.001, ****P <0.0001.

**Figure S5.**
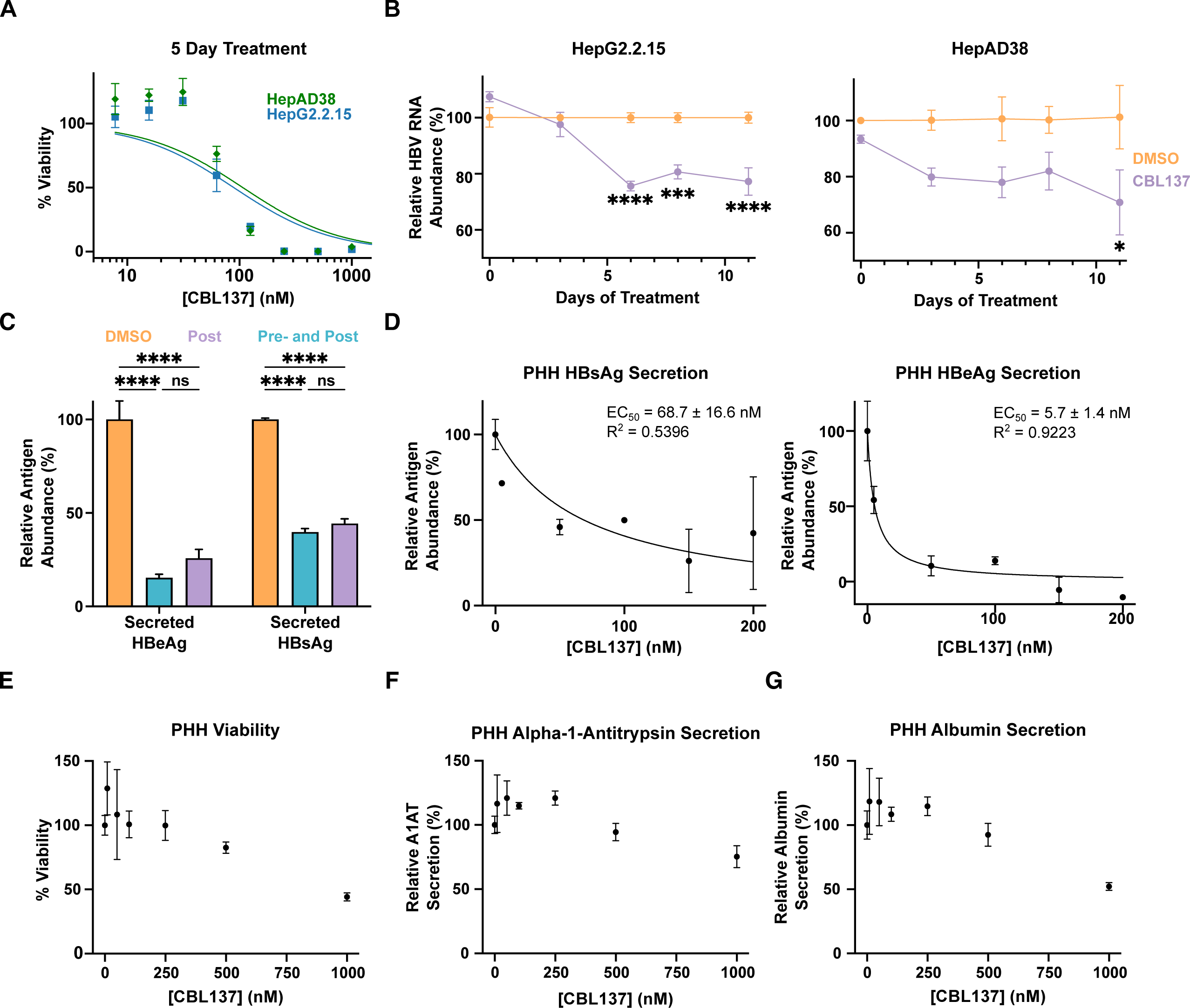
Impacts of long-term CBL137 treatment on HepG2.2.15 and HepAD38 cells, and primary human hepatocytes, related to Figure 5. (A) HepG2.2.15 and HepAD38 cells were cultured in media containing the indicated concentration of CBL137 for 5 days before viability was measured by CellTiter-Glo luminescence assay. (B) Quantification of relative total HBV RNA abundance in HepG2.2.15 (left) and HepAD38 (right) cells grown continuously in media containing either DMSO or 50 nM CBL137 at the indicated time points. (C) Quantification of HBsAg and HBeAg secretion from HepG2-NTCP cells that did or did not receive 24 hour pre-infection treatment with 125 nM CBL137 in addition to 5 day post-infection treatment. (D) Dose-dependent reduction of HBsAg and HBeAg secretion abundance in supernatant of primary human hepatocytes infected with HBV for 5 days and subsequently treated with the indicated doses of CBL137 for 2 days prior to analysis. (E) Dose-dependence of cell viability of primary human hepatocytes cultured and treated as in D. (F) Dose-dependence of alpha-1-antitrypsin secretion levels in the supernatant of primary human hepatocytes cultured and treated as in D. (G) Dose-dependence of albumin secretion levels in the supernatant of primary human hepatocytes cultured and treated as in D. Data represent means of 3 replicates ± SD and were analyzed by 2-way ANOVA with Šidak’s multiple comparison test (B) or Dunnett’s multiple comparisons test (E-G). *P < 0.05, **P < 0.01, ***P < 0.001, ****P <0.0001.

## STAR Methods

### RESOURCE AVAILABILITY

#### Lead contact

Further information and requests for resources and reagents should be directed to and will be fulfilled by the lead contact, Yael David.

#### Materials availability

All unique reagents generated in this study are available from the lead contact upon request and with a completed Material Transfer Agreement.

#### Data and code availability

- All sequencing data have been deposited at the Gene Expression Omnibus and are publicly available. Accession numbers are listed in the key resources table.
- This paper does not report original code.
- Any additional information required to reanalyze the data reported in this paper is available from the lead contact upon request.

### EXPERIMENTAL MODEL AND SUBJECT DETAILS

#### Mammalian cell lines and culture conditions

HEK293T, HepG2, HepG2-NTCP, HepG2.2.15, and HepAD38 cells were cultured in Dulbecco’s modified Eagle medium (DMEM) supplemented with 2 mM L-glutamine, 500 U/mL each of penicillin and streptomycin, and 10 % FBS and maintained in a humidified incubator at 37 °C with 5 % CO2. HepG2, HepG2-NTCP, HepG2.2.15, and HepAD38 cells were cultured in plates pre-treated with collagen (50 mg/mL working concentration, minimum 1 hr treatment). HepAD38 cells were additionally supplemented with tetracycline (0.5 µg/mL) for standard culturing but allowed to grow in media without tetracycline for two weeks prior to use for studies of HBV. Transfections of HEK293T cells were performed using Lipofectamine 2000 reagent according to manufacturer protocol. Transfections of HepG2 cells were performed with TransfeX Transfection Reagent (ATCC) according to manufacturer protocol. All cell lines were authenticated by the supplier and tested negative for mycoplasma contamination.

#### Bacterial strains used

*Escherichia coli* DH5α cells (Invitrogen) were used for all molecular cloning and large-scale plasmid production steps. *E. coli* BL21 cells (Invitrogen) were used for overexpression of human histone proteins and purification from inclusion bodies. All growth and expression conditions are outlined in method details.

## METHOD DETAILS

### Plasmid cloning

A single copy of the HBV genome (genotype D subtype ayw) was inserted into an empty pWM530 vector by restriction cloning. All restriction enzymes were purchased from New England Biolabs (NEB) and digestion reactions performed according to manufacturer protocols. A plasmid harboring a 1.1#x00D7; overlength genome of HBV (pHBV1.1X^82^, a gift from Prof. Stephan Urban, University Hospital Heidelberg, Heidelberg, Germany) was first digested with BspHI and EcoRI. The desired HBV genome fragment containing nt 1920-3182 was resolved by electrophoresis on a 1% agarose gel run at 130 V for 30 minutes at room temperature in 1X TAE buffer, cut out of the gel, and extracted and purified using a Qiaquick Gel Extraction Kit (Qiagen). The purified fragment was added to pWM530 digested with the same enzymes and ligated using T4 DNA ligase (NEB) according to manufacturer protocol. In parallel, the remaining portion of the HBV genome from upstream of nt 1 through the PsiI restriction motif was amplified from pHBV1.1X using PCR to introduce a 5’ BspHI restriction site. Both the PCR amplicon and the pWM530 vector containing a partial HBV genome were subsequently digested with BspHI and PsiI and ligated together to yield the plasmid pWM530-HBV1X. The pWM530-18#x00D7;601 construct was similarly prepared via two iterative instances of restriction enzyme digest and ligation to add 3 tandem 601 repeats onto the 3’end of a 12#x00D7;601 array in the pWM530 vector. The pWM530-X-601 chimeric construct was generated by first PCR amplifying the desired region of the HBV genome from the X promoter through the end of the X gene body using primers that introduced 5’ BglII and 3’ NheI restriction sites. Next the PCR amplicon and recipient pWM530-12#x00D7;601 plasmid were sequentially digested with NheI and either BglII (insert) or BamHI (vector). Resulting digested fragments were ligated with T4 ligase, transformed into DH5α cells, and the resulting plasmid sequence verified. All primer sequences used are reported in Table S2.

### DNA preparation for chromatin assembly

Mononucleosome-sized 601 DNA was prepared by PCR amplification of a DNA template containing one copy of the Widom 601 DNA sequence. PCR products were then pooled and purified with a QIAquick PCR Purification Kit (Qiagen), using water to elute from the final columns. Eluents were pooled, frozen, and lyophilized before being resuspended in buffer TE (10 mM Tris-HCl pH 7.6, 0.1 mM EDTA), quantified by NanoDrop OneC, and adjusted to a final concentration of approximately 1-1.5 g/L.

DNA templates used for chromatin fibers were prepared essentially as described before.^83^ *E. coli* DH5α cells were transformed with the desired pWM530 vector and used to inoculate 6 L of luria broth under ampicillin selection and grown at 37 °C for 18-24 hrs. Cultures were harvested and DNA purified using a Plasmid Giga Kit (Qiagen). Purified DNA was resuspended in buffer TE and adjusted to a concentration of ∼1 g/L. Plasmid was then digested with EcoRI and EcoRV (NEB) overnight to generate linear chromatin templates (i.e., dslDNA or Lin601^18^). Complete digestion was determined by agarose gel electrophoresis in 1X TAE buffer. Next, digestion reactions were fractionated by PEG 6000-induced precipitation as described previously to separate the ∼3.2 kb desired chromatin substrates from the 200-300 bp plasmid backbone fragments.^84^ Precipitated DNA was next resuspended in buffer TE, purified by phenol/chloroform extraction, and precipitated with absolute ethanol as described.^85^ DNA was finally resuspended in buffer TE, quantified by NanoDrop OneC, and adjusted to a concentration of 1-1.5 g/L. Buffer DNA was likewise prepared by digestion of a similar construct containing 8 repeats of the 155 bp mouse mammary tumor virus (MMTV) weak nucleosome positioning sequence with EcoRV, PEG precipitation, and phenol/chloroform extraction.

To generate DNA minicircles (i.e., cccDNA and Cir601^18^), the purified corresponding linear DNA was subject to intramolecular ligation conditions essentially as described previously.^25^ Ligations were performed at 16 °C with the following components: DNA starting at a concentration of 2 ng/µL, T4 Ligase (NEB) at 5 U/µL, and PEG 8000 at 5 % (v/v). Every 2-3 hrs, additional DNA was spiked into the reaction mixture to increase DNA concentration by 2 ng/µL. Ligations were allowed to proceed for 48-72 hrs before DNA was precipitated with PEG 6000 and purified by phenol/chloroform extraction as described above. Samples were then incubated overnight with Exonuclease III (NEB) to eliminate unligated species. As a final measure to confirm minicircle integrity, samples were also incubated for 10 minutes at 85 °C, allowed to cool, and resolved on 1% agarose gel. Subsequent digestion of DNA minicircles by EcoRI produced the original linear starting material.

### Histone expression and purification

Recombinant human histone proteins (H2A, H2B, H3.2, and H4) were purified as described before.^83^ Briefly, *E. coli* BL21 cells were transformed with pET21a expression plasmids containing each of the core histones and grown at 37°C in luria broth supplemented with 100 µg/mL ampicillin. After OD600 reached ∼0.6-0.8, isopropyl β-D-1-thiogalactopyranoside (IPTG) was added to cultures to a final concentration of 0.5 mM to induce protein expression. Cultures were harvested 4 hrs after induction and flash-frozen in liquid nitrogen.

Bacterial pellets were thawed in 1X PBS supplemented with 1 mM phenylmethylsulfonyl fluoride (PMSF) and subsequently lysed by rod sonication using a Branson Digital Sonifier with amplitude 40 %, 5 s on, 10 s off, for a total sonication time of 90 s. Lysates were cleared by ultracentrifugation at 30,000 xg for 30 minutes at 4 °C. Supernatant was discarded, and inclusion bodies were next resuspended in extraction buffer (1X PBS supplemented with 6 M guanidine hydrochloride). Pellets were allowed to extract at 4 °C overnight with agitation before again being subject to ultracentrifugation at 30,000 xg for 30 minutes. The supernatants were collected and histones purified by flash high-pressure reverse-phase liquid chromatography on an Agilent 1200 series instrument with semi-preparative Agilent 18C column (12 μm, 10 mm × 250 mm) using water supplemented with 0.1 % trifluoroacetic acid (TFA) as solvent A and a mixture of 90 % acetonitrile, 9.9 % water, and 0.1 % TFA as solvent B for the mobile phases. Purifications ran on a gradient from 0-70 % solvent A to B over 30 minutes with a flow rate of 4 mL/min. Samples were then analyzed by HPLC coupled to electrospray ionization MS (HPLC-ESI-MS) using an Agilent C18 column (5 μm, 4 × 150 mm) and an Agilent 6120 Quadrupole LC/MS spectrometer using the same solvent systems and gradient at a flow rate of 0.5 mL/min. Purified histones were lastly lyophilized and stored as a powder at -70 °C.

### Histone octamer assembly

To assemble histone octamers, monomeric core histones were resuspended in unfolding buffer (20 mM Tris-HCl pH 7.6, 6 M guanidine hydrochloride, 1 mM DTT) to a minimal concentration of 1 mg/mL and quantified. Core histones were next combined at the following stoichiometries to generate a slight excess of H2A/H2B dimer to assist with subsequent purification: 1.05:1.05:1:1 H2A:H2B:H3:H4. Total protein concentration of the mixture was adjusted to 1 g/L, and samples were dialyzed against octamer refolding buffer (10 mM Tris-HCl pH 7.6, 2 M NaCl, 1 mM EDTA, 1 mM DTT) at 4 °C using 3.5K MWCO Slide-A-Lyzer dialysis cassettes. A total of three rounds of dialysis against refolding buffer were performed, with the first exchange going overnight and the subsequent two lasting at least 6 hrs.

Following the final dialysis, samples were harvested, visible precipitate was cleared by centrifugation, and samples then concentrated using 30K MWCO Amicon centrifugal filter units. Finally, samples were centrifuged at >17,000 xg for a minimum of 10 minutes at 4 °C before being injected onto an AKTA 25L FPLC instrument and resolved over a SuperDex 200 10/300 Increase column, using octamer refolding buffer for the liquid phase. Fractions were analyzed by SDS-PAGE on 12 % acrylamide gels, and those containing intact octamers were pooled and concentrated using Amicon 30K MWCO centrifugal filter unites. Octamers were quantified, adjusted to 50 % (v/v) glycerol, and stored at -20 °C for future use.

### Chromatin reconstitution

Nucleosome core particles (NCPs), linear nucleosome arrays, and circular nucleosome arrays were all prepared by the same method of salt gradient dialysis with all steps at 4 °C. Substrate DNA and recombinant histone octamers were combined in approximately equimolar quantities (with respect to the expected nucleosome load of the DNA, i.e., 1 for mononucleosomal DNA, or 18 for HBV or 18#x00D7;601 DNAs) with final buffer conditions identical to octamer refolding buffer – 10 mM Tris-HCl pH 7.6, 2 M NaCl, 0.1 mM EDTA, 1 mM DTT. For nucleosome array assemblies MMTV DNA was also added, but at a lower stoichiometry to act as a buffer that facilitates array assembly without competing for octamer occupancy (0.2:1 MMTV:NCP). Optimal assembly stoichiometries were determined empirically for all templates, with Octamer:DNA stoichiometries being approximately 1.2:1 for mononucleosomes, 1.6:1 for 601 nucleosome arrays, and 1.8:1 for HBV nucleosome arrays. For differential saturation series of cccDNA, a ratio of 1.5:1 was used for undersaturated arrays, 1.7:1 was used for intermediate saturation, and 1.9:1 was used to achieve fully saturated arrays.

Chromatin assembly reactions were combined and mixed by gentle pipetting, centrifuged for 5 min at >17,000 xg at 4 °C, and then added to Slide-A-Lyzer Mini dialysis buttons pre-moistened in array initial buffer (10 mM Tris-HCl pH7.6, 1.4 M NaCl, 0.1 mM EDTA, 1 mM DTT) and dialyzed against 200 mL of the same buffer for 1 hr. Next, a peristaltic pump was used to transfer 350 mL of dilution buffer (10 mM Tris-HCl pH 7.6, 10 mM NaCl, 0.1 mM EDTA, 1 mM DTT) to the samples in initial buffer with a flow rate of 0.5-1.0 mL/min. After all dilution buffer was transferred, arrays were left to dialyze for at least 1 hr or up to overnight. Samples were next moved to a fresh 350 mL of dilution buffer and dialyzed for 6 hrs. Lastly, samples were moved into 300 mL of fresh dilution buffer and dialyzed for 1-2 hrs before harvesting.

Mononucleosome samples were simply pipetted out of dialysis buttons, centrifuged for 5 min at 17,000 xg, and supernatant (in case any precipitation was present) was quantified, subject to EMSA to validate assembly efficiency, and stored at 4 °C for up to 2 months. Chromatin array samples were similarly harvested to yield a mixture of assembled chromatin fibers, MMTV mononucleosomes, and free MMTV DNA. To separate the chromatin fibers, an equal volume of precipitation buffer (10 mM Tris-HCl pH 7.6, 10 mM NaCl, 10 mM MgCl2) was added to samples followed by a 20-minute incubation on ice. Next, samples were centrifuged for 10 minutes at 17,000 xg. Supernatant was gently pipetted off samples to avoid disturbing the barely visible pellets, to which a desired volume of array dilution buffer was next added. Pellets were left on ice for 10 minutes undisturbed to allow for the gradual resuspension of chromatin pellets before being quantified. Finally, EMSA was used to validate assemblies and harvests. Chromatin arrays were stored at 4 °C for up to one week before use.

Chaperone-assisted chromatin assembly was carried out essentially according to the method of An and Roeder.^30^ Briefly, 0.8 µL of 0.5 M NaCl, 6 µL of 2 µM yeast Nap1 (a kind gift from Shixin Liu), and 0.5 µL of 10 µM histone octamer were combined and gently mixed together, and subsequently diluted with 27 µL of HEG buffer (25 mM HEPES ph 7.6, 0.1 mM EDTA, 10% glycerol) before incubating on ice for 30 min. For reactions with ACF added, 0.4 µL of recombinant ACF (Active Motif) and 10X ATP-MgCl2 mix (final concentration of 5 mM each) were added, and all assembly reactions were then incubated at 27 °C for 3-4 hrs. Assembled chromatin arrays were subsequently harvested as described above for salt-gradient dialysis.

### Quantification of HBV genomic species

Viral DNA was extracted from the media using GeneJET Viral DNA/RNA Purification Kit following the manufacturer’s instructions (Thermo Fisher Scientific, USA) and quantified by TaqMan qPCR (PrimeTime Gene Expression Master Mix, IDT). A plasmid, harboring a 1.1#x00D7; overlength genome of HBV (gift from Prof. Stephan Urban, University Hospital Heidelberg, Heidelberg, Germany) was used to generate a calibration curve and determine samples genome equivalent (Geq).

To isolate intracellular viral transcripts, infected/transfected cells were lysed using TRIzol Reagent (Thermo Fisher Scientific, USA) and processed using Direct-zol DNA/RNA isolation kit (Zymo research, USA). RNA fraction was further treated with TURBO DNase (Thermo Fisher Scientific, USA) and purified using RNA clean & concentrator kit (Zymo research, USA). cDNA was synthesized using LunaScript RT SuperMix Kit (New England Biolabs, USA). SYBR green assay was used to determine HBV pregenomic expression (Luna Universal qPCR Master Mix, New England Biolabs, USA).

To quantify HBV cccDNA levels DNA isolated fraction from the infected cells was treated for 1h with T5 exonuclease^86^ (5U, New England Biolabs) and purified using a DNA clean and concentrator kit (Zymo research, USA). SYBR green assay was used for cccDNA detection with a calibration curve as described above to calculate HBV Geq. All primer sequences are presented in Table S2.

### Electrophoretic mobility shift assays (EMSAs)

Mononucleosome EMSAs were performed using polyacrylamide gel electrophoresis in 5 % acrylamide, 0.5 X TBE gels. A solution of 1 M sucrose was used as a loading buffer (1:3 dilution) for nucleosome samples. Gels were run for 30-40 minutes at a constant voltage of 130 V, stained with SYBR Gold dye diluted 1:10,000 in 0.5 X TBE buffer for 5-10 minutes, and DNA migration was visualized on an Amersham AI600 imager (GE/Cytiva) using the UV 312 nm channel. If needed, protein migration was visualized after DNA imaging by staining gels with Imperial Protein Stain and visualized on the AI600 instrument in the colorimetric channel.

Chromatin array EMSAs were performed similarly, except with a different gel formulation. Agarose-polyacrylamide gel electrophoresis (APAGE) gels were cast using the Mini-PROTEAN Tetra Handcast system (Bio-Rad). Before combining gel-casting reagents, the gel-facing sides of the 1.5 mm spacer plates, short plates, and 10-well gel combs were lubricated with a thin layer of 50 % (v/v) glycerol. Next, ultrapure water, 50X TAE (sufficient for a final concentration of 0.5 X), and powdered agarose (sufficient for a final concentration of 1 % w/v) were heated until dissolved. This solution was rapidly combined with 40 % acrylamide (37.5:1 mono:bis) solution (Bio-Rad) (sufficient for a final concentration of 2 %), APS (sufficient for 0.125 %), and TEMED (sufficient for 0.04 %), and added into the assembled gel casting cassettes. Gels were allowed to cool and polymerize for at least 1 hr. Once cool, gels were pre-run at 4 °C in 0.5 X TAE buffer for 3 hrs at 100 V. Next, chromatin samples could be loaded onto gels using sucrose loading solution and the gels run for approximately 1 hr at 120 V at room temperature before visualization as mononucleosome EMSAs.

### Atomic force microscopy

DNA-protein complexes were imaged using an Asylum MFP 3D Bio AFM (Oxford Instruments, Goleta CA) with an Olympus AC240TS probe in tapping mode at room temperature. The samples were diluted a suitable concentration (0.5-1.0 ng/µL DNA) in TEN100 (10 mM Tris-HCl pH 7.6, 0.1 mM EDTA, 100 mM NaCl), then 40 uL of prepared samples were slowly deposited to a freshly cleaved AP-mica for 5 minutes and rinsed with 1 mL ultrapure deionized water twice before being gently dried with UHP argon gas.

AFM images were collected at a speed of 0.5-1 Hz at 512 × 512-pixel resolution, with an image size of 2 μm. For analysis, raw images were exported into 8-bit grayscale Tiff images using the Asylum Research’s Igor Pro software and imported into FIJI/ImageJ (NIH) for detection of single particles and quantification of volume, surface area, and volume/surface area ratio using as has been done previously for studies of chromatin compaction via AFM.^87^ In order to assess single chromatin particles, rather than potential clusters of multiple fibers or residual MMTV mononucleosomes, only particles with volumes measuring between 2,000 and 15,000 nm^3^ were included in analyses.

### *In vitro* MNase-seq

Titration of micrococcal nuclease (MNase) was performed to determine enzyme concentrations that yielded partial-, fully-, or over-digested chromatin arrays. Chromatinized DNA (100 ng) was digested using MNase (NEB) in 25 µL reactions with 2.5–25 units of enzymes that was incubated for 5 minutes at 37 °C before 1 mM of CaCl2 was added and incubation continued for another 5 minutes at 37 °C. The reaction was stopped by adding 20 mM EDTA and 1 % SDS. DNA was subsequently purified using Clean and Concentrated-5 DNA columns (Zymo). Sequencing libraries were prepared using NEBNext Ultra II DNA Library Prep Kit – DNA repair, End Prep, USER, and Ligation of Unique Dual Index Barcodes or Unique Dual Index UMI Adaptors (NEB). Barcoding PCR was consistently carried out using only 5 PCR cycles with NEBNext High-Fidelity 2X PCR Mix and subsequent size selection using agarose gel or beads (Ampure XP). Libraries were sequenced on Illumina NextSeq500 or NextSeq1000 sequencers. To map to the HBV genome, paired-end reads were trimmed using a custom script and aligned with Bowtie2 v2.3.4.3 to the HBV genome (NC_003977.2).^88^ To account for any alignment fall off at the ends of the linear chromosome, NC003977.2 genome was mapped to both the EcoRI linearization start/end site or adjusted by 500 bp to match coordinates from Tropberger *et al*.^19^ The coordinates displayed throughout the figures are those that match the Tropberger *et al*.^19^ Because of the small genome size and the highly overlapping mononucleosome coverage, duplicates were not removed as this process removed many non-unique reads that were from bona fide original fragments. To verify this assumption, some experimental replicates used UMI barcodes and were processed using a custom script for UMI identification and duplicate removal (unix and R v4.1.0). For comparison across libraries, each sample was first subsampled to equal number of fragments (5 x 10^5^ read pairs) and NucleoATAC v0.3.4 was used to calculate fragment length distributions (pyatac sizes --not_atac) and mononucleosome fragment-centered coverage (pyatac cov --lower 140 --upper 200 -- not_atac --scale 1 --window 50).^89^ Mononucleosome coverage was smoothed with a 50 bp flat window to show peaks of nucleosome-sized fragment coverage that may be closer than the typical minimum ∼150 bp nucleosome spacing, which could indicate likely heterogeneity in the underlying population. The resulting bedgraph files were plotted in R using custom script or the Sushi package v1.30.0.^90^

### Enzyme-linked immunosorbent assays

HBV secreted antigen (HBeAg) and surface antigen (HBsAg) were quantitated by ELISA kit according to the manufacturer’s instructions (Autobio Diagnostics). Human albumin and alpha-1-antitrypsin were quantified by ELISA kit according to manufacturer instructions (Bethyl Laboratories). Signal was recorded using SpectraMax M5 plate reader (Molecular Devices).

### Immunostaining

Cells were fixed with 4% paraformaldehyde for 20 min and permeabilized using 0.3 % Tween-20 in PBS for 30 min at room temperature. Cells were subsequently blocked with MAXblock Blocking Medium (Active Motif) 1 hr at 37 °C, and then incubated with anti-HBcAg antibody (C1-5, Santa Cruz Biotechnology) diluted in MAXpack Immunostaining Media Kit (Active Motif) 1 hr at 37 °C. Nuclei were counterstained with Hoechst 33342 (Thermo Fisher) followed by extensive washes and imaging using EVOS FL microscope (Thermo Fisher Scientific).

### Magnesium-dependent self-association assay

Chromatin compaction was tested by magnesium-driven self-association as described previously.^28^ Briefly, a magnesium solution (100 mM MgCl2, 10 mM Tris-HCl pH 7.6, 10 mM NaCl) was titrated into the sample solution to raise the magnesium concentration in 0.5 mM increments. After each addition, samples were allowed to sit on ice for 10 min and then centrifuged for an additional 10 min at 17,000 xg at 4 °C. The concentration of soluble DNA from nucleosome arrays was measured by NanoDrop A260.

### *In vitro* transcription

All assays with the HeLaScribe Nuclear Extract *in vitro* Transcription System (Promega) were performed according to manufacturer protocols. For each transcription reaction, 150 ng of empty or chromatinized cccDNA was combined with 8 U HeLaScribe® Nuclear Extract, 1X transcription buffer, 3 mM MgCl2, and 500 µM each rATP, rCTP, rGTP, and rUTP. Reactions were incubated at 30 °C for 60 minutes. Next, to ensure that no template DNA carried over to contaminate downstream results, 1 µL of DNase I (Amplification Grade, Invitrogen) was added to the mixture and incubated for 20 minutes at 30 °C. Finally, HeLa Extract Stop Solution was added to the reactions. RNA was then purified from the quenched reaction mixtures using Monarch® RNA Cleanup Kit (NEB) according to manufacturer protocols and subsequently quantified by NanoDrop.

HepG2 nuclear extract was prepared by the method of Dignam and Roeder.^34^ Briefly, ∼1#x00D7;10^8^ frozen HepG2 cells were thawed and resuspended in 5 mL buffer A (10 mM HEPES pH 7.9, 1.5 mM MgCl2, 10 mM KCl, 1 mM DTT) and incubated on ice for 10 min. Cell suspension was then centrifuged at 400 x g for 10 min. at 4 °C and supernatant was subsequently discarded. The cell pellet was resuspended in 2 mL of buffer A, transferred to a pre-chilled glass Dounce homogenizer, and lysed with 10 gentle strokes of a loose pestle. Lysate was subsequently centrifuged at 400 x g for 10 min. at 4 °C to pellet nuclei, after which the supernatant was removed. Pellets were then spun again at 17,000 x g for 20 min. at 4 °C, and supernatant was removed to yield crude nuclei. Nuclei were then resuspended in 2 mL buffer C (20 mM HEPES pH 7.9, 25 % (v/v) glycerol, 420 mM NaCl, 1.5 mM MgCl2, 1 mM PMSF, 1 mM DTT), transferred to a pre-chilled Dounce homogenizer, and lysed with 10 strokes of a tight pestle. The resulting homogenate was incubated at 4 °C for 30 minutes with gentle spinning with a magnetic stir bar for 30 min. Next, homogenate was centrifuged at 4 C for 30 min at 17,000 x g. The resulting clarified supernatant was next dialyzed against 200 mL of buffer D (20 mM HEPES pH 7.9, 20 % (v/v) glycerol, 100 mM KCl, 0.2 mM EDTA, 1 mM PMSF, 1 mM DTT) for 5 hrs at 4 °C. Finally, dialysate was centrifuged at 4 °C for 20 min. at 17,000 x g to pellet any undesired precipitate and supernatant was removed, quantified, aliquoted, and flash frozen in liquid nitrogen. Extracts were validated for transcription using the positive control reagents provided in the commercial HeLaScribe (Promega) kit and subsequent *in vitro* transcription assays with HepG2 extract were performed as described above for the commercial HeLa extract, using an equivalent amount of HepG2 extract to match the activity of the commercial reagent.

RNA samples were converted to cDNA using High-Capacity RNA-to-cDNA Kit (Applied Biosystems). Quantitative PCR reactions were performed using the iTaq Universal SYBR Green Supermix (Bio-Rad) with each 10 µL reaction containing 5 ng cDNA and 500 nM of both the forward and reverse primers for the reaction. Relative gene expression levels (as in drug treatment experiments) were calculated using the ΔΔCT method. Absolute RNA quantification was calculated using a standard curve as described above for total viral RNA, and as described by D’Arienzo, et al.^35^, for transcript-specific RT-qPCR. Briefly, the HBVqT1F/R primer pair was used to determine the abundance of pgRNA/preC transcripts. The total transcript number for these species was then subtracted from that of the HBVqT2F/R primer pair to determine the abundance of preS1 transcript species, as this primer pair covers a region present in both pgRNA/preC and preS1 transcripts. Likewise, the total number of preS1 transcripts was subtracted from that of the HBVqT3F/R primer pair to determine the abundance of preS2 transcripts, and so on with the HBVqT4F/R primer pair to detect X transcripts. All primer sequences are presented in Table S2.

### Immunoblotting

All samples were resolved on 4-20% Mini-Protean TGX precast protein gels (Bio-Rad) and transferred to a PVDF membrane for western blot analysis. Membranes were blocked with 5 % BSA (w/v) in PBS-T (1 X PBS supplemented with 0.1 % Tween-20) for 30 minutes at room temperature. Membranes were next rinsed three times each with PBS-T for 5 minutes, and incubated with primary antibodies diluted in PBS-T at room temperature for 1-2 hours. Membranes were then washed with PBS-T as before and then incubated with secondary antibodies for one hour at room temperature. Finally, membranes were washed as before and imaged in the 680 nm and 800 nm channels using an Odyssey CLx imager (LI-COR). All primary antibodies were used in 1:1,000 dilutions, and all secondary antibodies were used in 1:15,000 dilutions.

### Full-length 5’ RACE

Cellular RNA was isolated using the RNeasy Plus Mini kit (Qiagen), and *in vitro* transcribed RNA was purified using the using the Monarch® RNA Cleanup Kit (NEB). Full-length 5’ RACE was performed using the GeneRacer™ with Superscript™ III RT kit (ThermoFisher Scientific) as described in the manual with modifications adapted from the methods of Stadelmeyer *et al*.^17^ The 5’RACE PCR was ran using the Platinum™ SuperFiTM DNA Polymerase (ThermoFisher Scientific), with touchdown PCR according to the following steps: 1#x00D7; 98 °C 3 min; 5#x00D7; (98 °C 10 s; 72 °C 3 min); 5#x00D7; (98 °C 10 s; 70 °C 3 min); 25#x00D7; (98 °C 10 s; 64.4 °C 20s; 72 °C 3 min); 72 °C 10 min. PCR products were subsequently resolved on a 1% agarose gel and visualized with ethidium bromide.

### Peptide synthesis

Solid-phase peptide synthesis using Fmoc chemistry was performed using a Biotage Initiator+ Alstra automated peptide synthesizer following manual coupling of the first (C-terminal) amino acid as described elsewhere.^91^ Coupling reagents and Fmoc-protected amino acids were purchased from AGTC Bioproducts, and all other reagents from Sigma Aldrich. Briefly, Rink Amide (ChemMatrix) resin was swelled in DMF and deprotected with 20 % piperidine in DMF for 1 hr. Then 5 molar eq. of Fmoc-protected amino acid dissolved in 0.5 M HBTU and 0.5 M HOBt in DMF was added to resin along with 10 molar eq. DIEA. The coupling reaction was allowed to proceed at room temperature for 20 min. with agitation by N2 bubbling. After coupling, unreacted resin was capped with acetic anhydride and DIEA, and resin was moved to the automated synthesizer for peptide elongation using a similar protocol. Following completion of the synthesis, peptide was cleaved from resin by incubation with a cocktail of 95 % TFA, 2.5 % TIS, and 2.5 % water. Crude cleavage mixture was purified by RP HPLC using an Agilent C18 column with a 0-70 % gradient from Buffer A (Water + 0.1 % TFA) to Buffer B (90 % Acetonitrile, 9.9 % water, 0.1 % TFA). Fractions containing the desired peptide were pooled, quantified, and lyophilized prior to storage at -80 °C.

### *In vivo* MNase-seq

HEK293T cells were transfected with recombinant cccDNA 4 or 24 hrs prior to harvest for MNase-seq. The nuclei of HEK293, HepAD38, and HepG2.2.15 cells were purified by incubating cells in Nuclei isolation buffer for 10 minutes (0.1 % Igepal, 0.1 % Triton X -100, 1 mM DTT in 1 X PBS), mechanically lysing with 10 strokes of loose (A) pestle Dounce homogenizer, and washing nuclei twice in 10 mL of Wash Buffer with Tween (10 mM Tris-HCl pH 7.5, 2.5 mM NaCl, 3 mM MgCl2, and 0.1 % Tween-20). Titration of MNase was similarly performed to determine enzyme concentrations that yielded partial-, fully-, or over-digested genomic DNA. MNase (NEB) was added to 2 x 10^6^ nuclei in 50 µL reaction volume with 250– 2000 units of enzyme that was incubated for 5 minutes at 37 °C before 1 mM of CaCl2 was added and incubation continued for another 5 minutes at 37 °C. The reaction was stopped by adding 20 mM EDTA and 1 % SDS. DNA was subsequently purified using Clean and Concentrated-5 DNA columns (Zymo).

Sequencing libraries were prepared using NEBNext Ultra II DNA Library Prep Kit – DNA repair, End Prep, USER, and Ligation of Unique Dual Index UMI Adaptors (NEB). Barcoding PCR was consistently carried out with NEBNext High-Fidelity 2X PCR Mix and subsequent size selection using agarose gel or beads (Ampure XP). Libraries were then pooled and enriched for HBV-containing reads using custom-designed xGen Lockdown probes (Integrated DNA Technology) of 60 bp each tiling the entire HBV genome and oligo capture protocol was followed essentially per the manufacture’s protocol, except that the probe annealing steps were performed at 50 °C (Table S3 for probe sequences). Enriched libraries were subsequently amplified using 11 PCR cycles with NEBNext High-Fidelity 2X PCR Mix and purified using Ampure XP beads. Libraries were sequenced on Illumina NextSeq500 or NextSeq1000 sequencers with paired-end 50 bp x 50 bp or 100 bp x 100 bp reads. Raw data was processed by removing duplicates using custom UMI identifier script and then reads were aligned using Bowtie2 v2.3.4.3 to the HBV genome (NC_003977.2).^88^ To account for any alignment fall off at the ends of the linear chromosome, NC003977.2 genome was mapped to both the EcoRI linearization start/end site or adjusted by 500 bp to match coordinates from Tropberger *et al*.^19^ The coordinates displayed throughout the figures are those that match the Tropberger *et al*.^19^ NucleoATAC v0.3.4 was used to calculate fragment length distributions (pyatac sizes --not_atac) and mononucleosome fragment-centered coverage (pyatac cov --lower 140 --upper 200 --not_atac --scale 1 --window 50).^89^ Mononucleosome coverage was smoothed with a 50 bp flat window, as for *in vitro* MNase-seq. The resulting bedgraph files were plotted in R using custom script or Sushi package v1.30.0.^90^

### RNA sequencing

Samples for RNA-sequencing were harvested into 1 mL TRIzol Reagent (ThermoFisher) and phase separation was induced with 200 µL chloroform. RNA was extracted from 350 µL of the aqueous phase using the miRNeasy Mini Kit (Qiagen) on the QIAcube Connect (Qiagen) according to the manufacturer’s protocol. Samples were eluted in 38-42 µL RNase-free water. After RiboGreen quantification and quality control by Agilent BioAnalyzer, 500 ng of total RNA with RIN values of 9.9-10 underwent polyA selection and TruSeq library preparation according to instructions provided by Illumina (TruSeq Stranded mRNA LT Kit), with 8 cycles of PCR. Samples were barcoded and run on a NovaSeq 6000 in a PE100 run, using the NovaSeq 6000 S4 Reagent Kit (200 Cycles) (Illumina). An average of 28 million paired reads was generated per sample. Ribosomal reads represented at most 0.38 % of the total reads generated and the percent of mRNA bases averaged 87 %.

Initial assessment of RNA-seq data was done by pseudoalignment of reads (kallisto version v0.48.0) to hg38 (GRCh38.p13) and differential gene expression analysis of transcript abundances in R (DESeq2 v1.36.0) with the time points compared to the no transfection baseline (0 hr) using a Wald test with a = 0.05 and log2 fold-change shrinkage with a beta prior (DESeq(de.peaks, betaPrior=TRUE)).^94,95^ For each set of comparisons performed, lowly expressed genes were filtered out by removing those whose mean across all conditions were ≤ 10 tags per million. To fully align reads to the HBV genome, every nucleotide of which is transcribed, paired-end reads were trimmed with a custom script, aligned with Bowtie2 v2.3.4.3 to the HBV genome (NC003977.2),^88^ and duplicates were removed using Picard v2.20.3. The exact same alignment was also performed against hg38 for direct comparison of the number of reads that map to the human genome. To account for any alignment fall off at the ends of the linear chromosome, NC003977.2 genome was mapped to both the EcoRI linearization start/end site or adjusted by 500 bp to match coordinates from Tropberger *et al*.^19^ The coordinates displayed throughout the figures are those that match the Tropberger *et al*.^19^ Bedtools was used to calculate coverage along the HBV genome in a strand-specific manner (genomeCoverageBed -pc -strand +/-) and plots were made using custom script (R v4.1.0) or Sushi package v1.30.0.^90,96^

### Chromatin immunoprecipitation (ChIP)-sequencing

Freshly harvested cells were fixed with 1% formaldehyde for 10 min, after which the reaction was quenched by the addition of glycine to a final concentration of 0.125 M. Fixed cells were washed twice with PBS, snap frozen and store at -80 °C. Frozen cell pellets were sent to the MSKCC Epigenetics Research Innovation Lab for processing. Cells were resuspended in SDS buffer (100 mM NaCl, 50 mM Tris-HCl pH 8.0, 5 mM EDTA, 0.5 % SDS, 1#x00D7; protease inhibitor cocktail from Roche). The resulting nuclei were spun down, resuspended in the immunoprecipitation buffer at 1 mL per 0.5 million cells (SDS buffer and Triton Dilution buffer [100 mM NaCl, 100 mM Tris-HCl pH 8.0, 5 mM EDTA, 5% Triton X-100] mixed in 2:1 ratio with the addition of 1#x00D7; protease inhibitor cocktail) and processed on a Covaris E220 Focused-ultrasonicator to achieve an average fragment length of 200-300 bps with the following parameters: PIP=140, Duty Factor=5, CBP/Burst per sec=200, Time = 1200s. Chromatin concentrations were estimated using the Pierce™ BCA Protein Assay Kit (ThermoFisher Scientific) according to the manufacturer’s instructions. The immunoprecipitation reactions were set up in 500uL of the immunoprecipitation buffer in Protein LoBind tubes (Eppendorf) and pre-cleared with 50 µL of Protein G Dynabeads (ThermoFisher Scientific) for two hours at 4 °C. After pre-clearing, the samples were transferred into new Protein LoBind tubes and incubated overnight at 4 °C with the indicated antibodies (Key Resources Table). The next day, 50 µL of BSA-blocked Protein G Dynabeads were added to the reactions and incubated for 2 hours at 4 °C. The beads were then washed two times with low-salt washing buffer (150 mM NaCl, 1 % Triton X-100, 0.1 % SDS, 2 mM EDTA, 20 mM Tris-HCl pH 8.0), two times with high-salt washing buffer (500 mM NaCl, 1 % Triton X-100, 0.1 % SDS, 2 mM EDTA, 20 mM Tris-HCl pH 8.0), two times with LiCL wash buffer (250 mM LiCl, 10 mM Tris-HCl pH 8.0, 1 mM EDTA, 1 % Na-Deoxycholate, 1 % IGEPAL CA-630) and one time with TE buffer (10 mM Tris-HCl pH 8.0, 1 mM EDTA). The samples were then reverse-crosslinked overnight in elution buffer (1 % SDS, 0.1 M NaHCO3) and purified using the ChIP DNA Clean & Concentrator kit (Zymo Research) following manufacturer instructions. After quantification of the recovered DNA fragments, libraries were prepared using the ThruPLEX®DNA-Seq kit (Takara) following the manufacturer’s instructions, purified with SPRIselect magnetic beads (Beckman Coulter), and quantified using a Qubit Flex fluorometer (ThermoFisher Scientific) and profiled with a TapeStation (Agilent). The libraries were sequenced on an Illumina NovaSeq 6000 at 30-40 million 100 bp paired-end reads per library through the MSKCC Integrated Genomics Operation core facility.

ChIP sequencing reads were trimmed and filtered for quality and library adapters using version 0.4.5 of TrimGalore (https://www.bioinformatics.babraham.ac.uk/projects/trim_galore), with a quality setting of 15 and running cutadapt (v1.15) and FastQC (v0.11.5). Reads were aligned to human assembly hg38 and HBV assembly NC_003977.2 with version 2.3.4.1 of bowtie2 and were deduplicated using MarkDuplicates in version 2.16.0 of Picard Tools. Read density profiles were created using the BEDTools suite v2.29.2 for human and deepTools ‘bamCoverage’ v3.3.0 for HBV, normalized to 10 million uniquely mapped reads and with read pileups extended to the length of each paired fragment for HBV and extended 200 bp for human. To ascertain enriched regions in the human-aligned data, MACS2 was used with a p-value setting of 0.001 and run against a matched input control for each condition.^92^ Metagene profile composite plots were created for human alignments at gene bodies from Gencode version 28 using deepTools ‘plotHeatmap’ on normalized bigwigs with average signal sampled in 25 bp windows and flanking region defined by the surrounding 3 kb and scaled to 1 million uniquely mapped reads. For the HBV analysis, the genome was divided into 300 bp bins using BEDTools ‘makewindows’ and normalized signal was obtained by running ‘computeMatrix’ and ‘multiBigwigSummary’ in deepTools v3.3.0 using the NC_003977.2 bigwigs as input. The average difference in normalized read count from DMSO and CBL137 treated samples was then plotted in a heatmap and analyzed using GraphPad Prism. Read density profiles along the HBV genome were visualized using the integrative genomics viewer webtool.^93^

### Cell viability assays

Cells were seeded (10,000 cells/well) in the internal 60 wells of black, clear-bottom 96-well tissue culture plates (Corning). Wells along the perimeter of the plate were filled with PBS to minimize evaporation in the central wells. 24 hrs later, seeding media was aspirated off the adherent cells and replaced with media containing 0.1 % DMSO and a 2-fold dilution series of the indicated drug (or vehicle-only control). Cells were left to grow for 48 hrs of treatment (unless indicated otherwise) and then cytotoxicity was measured using the CellTiter-Glo Luminescent Cell Viability Assay (Promega) according to manufacturer protocol with a Molecular Devices Spectramax M3 plate reader.

### Thermal stability assays

Chromatin thermal stability was measured using a Protein Thermal Shift kit (Applied Biosystems). For each reaction, SYPRO Orange dye (Invitrogen, 10X final concentration), recombinant mononucleosomes/chromatin fibers (∼60 ng/µL DNA, final concentration, and DMSO or the indicated drug (final concentration 0.1 % DMSO) were combined and diluted with nucleosome dilution buffer (10 mM Tris-HCl pH 7.6, 100 mM NaCl, 0.1 mM EDTA, and 1 mM DTT) to a final volume of 10 µL per well in 384-well plates. Fluorescence melt curve data was acquired using a QuantStudio 5 Real-Time PCR System (Applied Biosystems) with the following method: initial ramp rate of 1.6 °C/s to 25 °C with a 5-minute hold time at 25 °C, followed by a second ramp rate of 0.05 °C/s to 99.9 °C with a 2-minute hold time at 99.9 °C. Melting temperatures were calculated using the Protein Thermal Shift software (Applied Biosystems) using the first derivative method.

### Generation of HBV stocks and HBV infection

HepG2.2.15 cells^42^ were cultured in DMEM with 10 % FBS and 200 mg/ml G418 (Sigma-Aldrich) until they reached a confluency of 100 %. Media was changed to DMEM-F12 supplemented with 10 % FBS, 1 % Pen/Strep. Media was filtered through a 0.22 μm filter (Millipore, Darmstadt, Germany), PEG-8000 was added (Sigma-Aldrich, final concentration of 6 %) and incubated overnight at 4°C. Virus was concentrated by centrifugation, 10,000 x g, 1 hr at 4°C and pelleted was resuspended using DMEM with 10 % FBS at 1/100 the original volume. Virus was aliquoted into cryovial tubes and cryopreserved at -80°C until use.

HepG2-NTCP cells^97^ (gift from Prof. Stephan Urban, University Hospital Heidelberg, Heidelberg, Germany) were maintained in DMEM supplemented with 10 % FBS, 1 % Pen/Strep and 5 µg/ml Puromycin (Goldbio). Cells were seeded into collagen-coated 24-well plates (50 µg/ml) in a density of 2×10^5^ cells/well and cultured in regular DMEM supplemented with 10 % FBS, 1 % Pen/Strep overnight. Cells were incubated with HBV inoculum diluted in media containing 4 % PEG-8000 (1000 Genome Equivalents [GEq]/mL). The next day, the virus inoculum was removed, and the cells were extensively washed with PBS and cultured in the presence of 2.5 % DMSO, with the addition of examined inhibitor or vehicle at the indicated concentration.

### Primary Human Hepatocytes

Cryopreserved Human Hepatocytes (Massachusetts General Hospital), were thawed at 37 °C, pelleted by centrifugation at 50 x g for 10 min and resuspended in hepatocyte culture medium, which consisted of 1#x00D7; DMEM, 10% FBS, Insulin-Transferrin-Sodium Selenite Supplement (Gibco,11360), Dexamethasone (Gibco 11040), Glucagon (Gibco 25030) and 1 % Pen/Strep. For experiments with HBV infection, hepatocytes were cocultured with NIH-3T3 J2 cell line stromal cells in PDMS microwells as previously described.^98^ Briefly, to generate 3D cell aggregates, PDMS microwells surface was first treated 5 % Pluronic F127 (Sigma Aldrich) for 1 hour at 37 °C to block absorption. Wells were extensively washed first with PBS and subsequently with DMEM. Next, hepatocytes were mixed in a 2:1 ratio with J2-3T3 cells in a 24 well format (120,000:60,000 per well) and the cell mixture was added to the wells, centrifuged at 50 x g for 10 min and incubated for 72 hours at 37 °C to allow aggregate formation. For viral infections, cells were incubated with HBV inoculum diluted in media (final concentration 300 Genome Equivalents [GEq]/mL) for 24 hours at 37 °C before the viral inoculum was removed, and the cells were extensively washed with fresh media. After cells had been infected for 5 days, they were treated with the indicated CBL137 concentrations (or vehicle-only control) for 48 hrs and HBsAg, HBeAg, viral RNA, and cccDNA were quantified as described above. To assess CBL137 impact on hepatocyte viability and function, freshly thawed viable hepatocytes were counted and plated in a density of 1#x00D7;10^6^ cells/mL on collagen coated 24-well plates. The following day, cells were washed and CBL137 (or vehicle-only control) was added to the media in the indicated concentrations. After 48 hrs of treatment, media was harvested for albumin and alpha-1-antitrypsin quantitation by ELISA and measurement of cell viability.

## QUANTIFICATION AND STATISTICAL ANALYSES

Statistical analyses were performed in GraphPad Prism v10.2.0. Specific tests used, error bar information, and number of experimental repeats can be found in figure legends. No statistical methods were used to predetermine sample sizes, and investigators were not blinded during experiments or analysis.

## Supplemental Item Titles

**Table S1.**
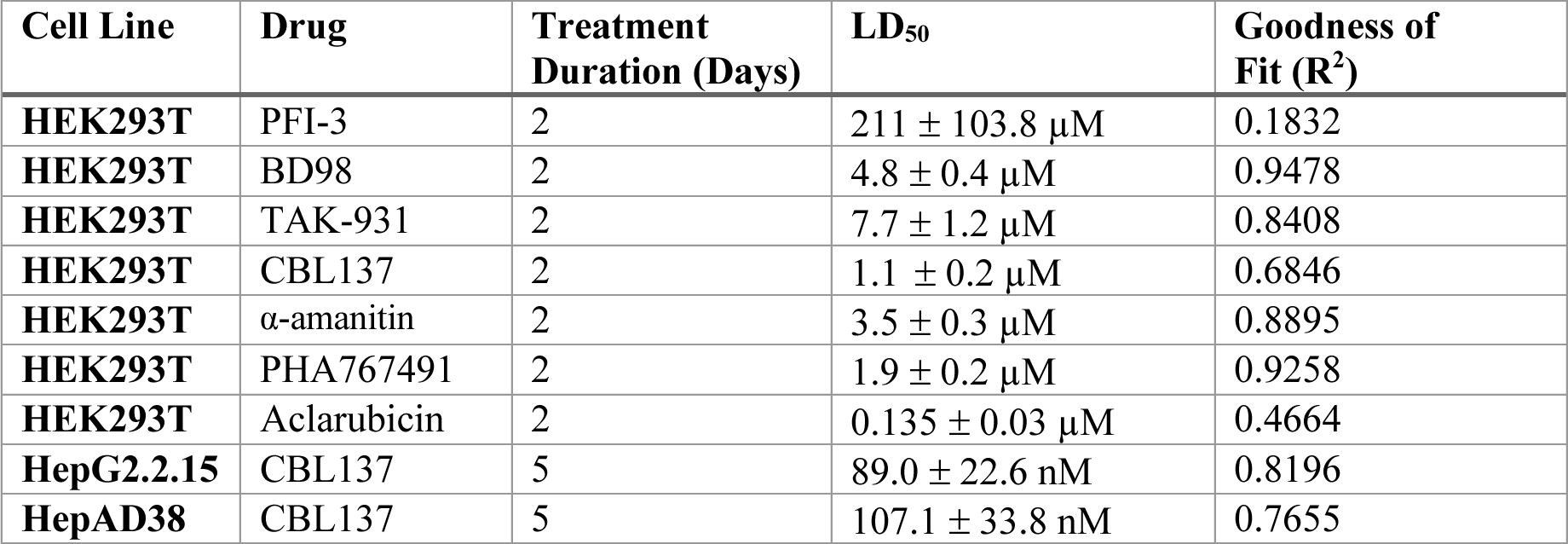
Cell viability calculations for tested drugs, related to Figures S4 and S5.

**Table S2.**
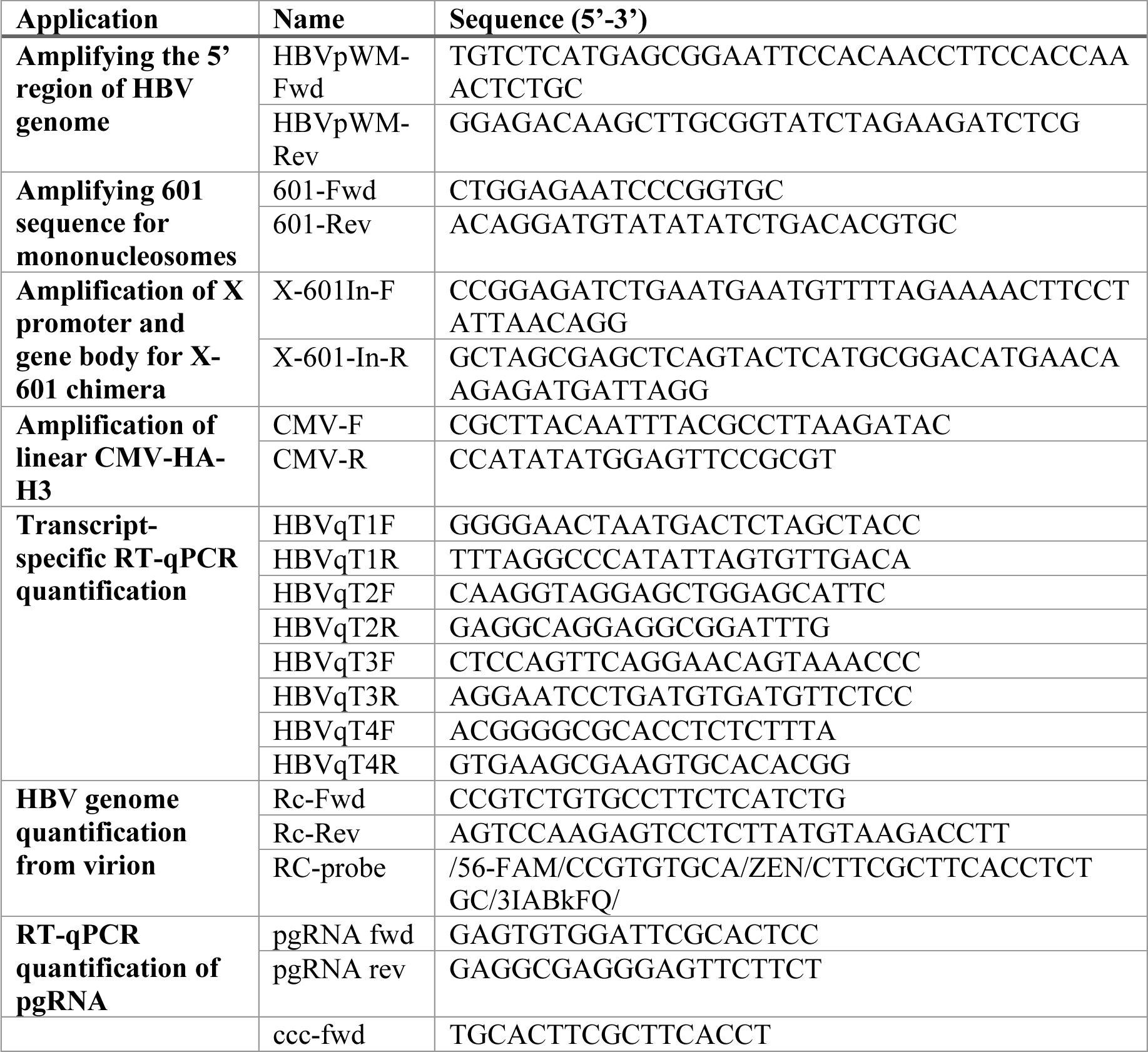
Primer sequences used for standard and quantitative PCR, related to STAR Methods.

**Table S3.**
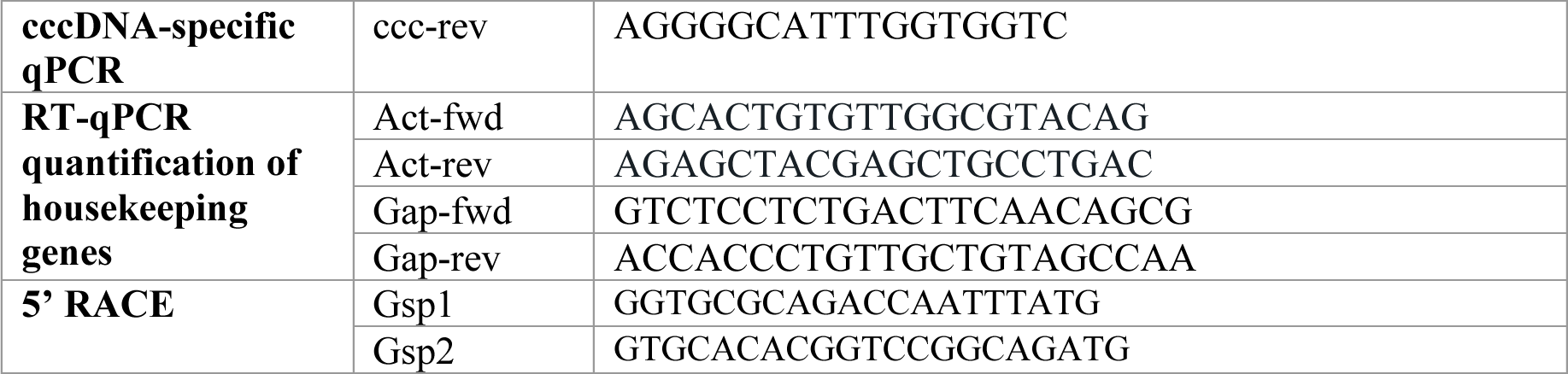
Probe sequences used for HBV enrichment in *in vivo* MNase-seq, related to STAR Methods.

